# Inhibitory KIRs decrease HLA class II-mediated protection in Type 1 Diabetes

**DOI:** 10.1101/2024.03.27.586933

**Authors:** Laura Mora Bitria, Bisrat J. Debebe, Kelly Miners, Kristin Ladell, Charandeep Kaur, James A. Traherne, Wei Jiang, Linda Hadcocks, Nicholas A.R. McQuibban, John Trowsdale, F Susan Wong, Nikolas Pontikos, Christoph Niederalt, Becca Asquith

## Abstract

Inhibitory killer cell immunoglobulin-like receptors (iKIRs) are a family of inhibitory receptors that are expressed by natural killer cells and late-stage differentiated T cells. There is accumulating evidence that iKIRs regulate T cell-mediated immunity. Recently, we reported that T cell-mediated control was enhanced by iKIRs in chronic viral infections. We hypothesized that in the context of autoimmunity, where an enhanced T cell response might be considered detrimental, iKIRs would have an opposite effect. We studied Type 1 diabetes (T1D) as a paradigmatic example of autoimmunity. In T1D, variation in the Human Leucocyte Antigen (HLA) genes explains up to 50% of the genetic risk, indicating that T cells have a major role in T1D etiopathogenesis. To investigate if iKIRs affect this T cell response we asked whether HLA associations were modified by iKIR genes. We conducted an immunogenetic analysis of a case-control T1D dataset (N= 11,961) and found that iKIR genes, in the presence of genes encoding their ligands, have a consistent and significant effect on protective HLA class II genetic associations. Our results were validated in an independent data set. We conclude that iKIRs significantly decrease HLA class II protective associations and suggest that iKIRs regulate CD4+ T cell responses in T1D.

## Introduction

Type 1 Diabetes (T1D) is a common autoimmune disease characterized by insulin-deficiency due to the destruction of insulin-producing islet β-cells. The exact aetiology of T1D remains elusive, but environmental triggers are thought to initiate the break in peripheral tolerance in genetically susceptible individuals^1^. The largest genetic contributors to susceptibility to T1D are the human leucocyte antigen (HLA) genes^2,3^. Within the HLA region, the closely linked classical class II *HLA-DRB1*, *HLA-DQB1* and *HLA-DQA1* genes display the strongest associations indicating that CD4+ T cells have a major role in T1D etiopathogenesis. In particular, *DRB1*04:01/02/04/05-DQA1*03:01-DQB1*03:02* and *DRB1*03:01-DQA1*05:01-DQB1*02:01* haplotypes are associated with the highest T1D susceptibility whereas *DRB1*15:01-DQA1*01:02-DQB1*06:02* is associated with dominant protection^4^.

Here we study a family of inhibitory receptors called inhibitory killer-cell immunoglobulin-like receptors (iKIRs). iKIRs are expressed predominantly on natural killer cells and, at a lower frequency, on late stage differentiated T cells. The ligands of iKIRs are HLA class I molecules which they bind in broad allele groups, e.g. KIR3DL1 binds HLA B alleles with a Bw4 motif at positions 77-83^5^. The iKIR genes and the genes encoding their HLA class I ligands are located on different chromosomes and so are inherited independently. Consequently, it is common for individuals to have one or more iKIRs without the corresponding ligand; if an individual is positive for a given iKIR as well the matching ligand we refer to that iKIR as “functional”. iKIRs play a major role in regulating innate NK-cell mediated responses but there is increasing evidence that iKIRs also modulate adaptive T cell responses^6-8^. In particular, iKIRs have been reported to increase activated T cell survival and to dampen effector function. The two main mechanisms of increased T cell survival are inhibition of activation induced cell death (attributed to iKIRs expressed on T cells) and inhibition of NK-killing of activated T cells (attributed to iKIRs expressed on NK cells)^9-11^. We have previously found that iKIRs together with their ligands significantly enhance CD8+ T cell survival in humans^8^. Furthermore, iKIRs with their ligands also enhance protective and detrimental HLA class I associations and have a significant impact on the clinical outcome of three different chronic viral infections^11,12^. We distinguish this modulation of protective and detrimental HLA associations by functional iKIR, which we suggest is due to a modulation of T cell responses by iKIR, from a main effect of functional iKIR, which we suggest is more likely to be NK-cell mediated^11,12^. Evidence for a main effect of iKIR or functional iKIR in T1D is weak. Since 2003, 15 studies have reported KIR gene associations with T1D risk, but none has been consistently reproduced and, in a recent metanalysis, no associations survived correction for multiple comparisons^13^. A few studies have explored functional iKIR associations i.e. associations between iKIR-HLA ligand gene pairs and T1D^14-16^ but again, results are not consistent across the studies. The impact of functional iKIR on HLA associations has not been studied.

We postulated that, given their impact on adaptive immune responses in chronic viral infections, iKIRs might play an analogous role in autoimmunity. Associations between HLA class II genes and T1D are clear evidence that CD4+ T cells play a role in T1D. We hypothesized that if iKIRs modulate CD4+ T cell responses then this should be manifest as an iKIR modulation of HLA class II genetic associations. We sought to test this hypothesis by investigating the impact of iKIRs on HLA class II disease associations in T1D using a large (N=11,961) case-control dataset from the UK Genetics Resource Investigating Diabetes (GRID). We identified a consistent and significant functional iKIR modification of HLA class II protective associations. The size of this effect was striking, for instance the odds ratio of iKIR in *DQA1*01:02-DQB1*06:02+* individuals is 6.12; one of the largest genetic effects reported in T1D in recent decades and is replicated across all protective class II genotypes. These findings are reproduced in a smaller independent dataset consisting of 339 US multiplex T1D families from the Human Biological Data Interchange. Our immunogenetic analyses show that genes encoding iKIRs with their ligands decrease protective class II genetic associations, consistent with a picture in which iKIRs modulate T cell-mediated regulation of autoimmunity.

## Results

We previously reported that functional iKIR genes (gene pairs encoding both the iKIR and its HLA class I ligand) enhanced protective and detrimental HLA disease associations in three chronic viral infections^11,12^. Here, we asked whether in the context of autoimmunity, where an enhanced T cell response might be considered detrimental, functional iKIRs had the opposite effect, i.e. that HLA associations were not enhanced but weakened by functional iKIR genes.

We chose T1D as a paradigmatic example to study the effect of iKIRs in autoimmunity. Our primary cohort was a T1D case-control cohort which consisted of (after removal due to missingness) of 6,219 cases and 5,742 controls (see **Materials and methods**). We first studied the impact of iKIRs on detrimental HLA class II disease associations. We found that most detrimental class II genotypes were not impacted by iKIRs (**Fig. S2**) and that the two detrimental genotypes which were modified by iKIRs were modified in opposite directions. Overall, there was no evidence of consistent iKIR modification (P=0.46). We next studied HLA protective class II associations starting with the best documented class II protective genotype: *DRB1*15:01-DQA1*01:02-DQB1*06:02*.

### iKIR score modifies the *DRB1*15:01-DQA1*01:02-DQB1*06:02* protective association

The *DRB1*15:01-DQA1*01:02-DQB1*06:02* compound genotype has repeatedly been described to confer protection from T1D^17^ and this protective effect is reproduced in our cohort (ln[OR]= -3.75, P=1.05x10^-157^). There is some evidence that the protection associated with the *DRB1*15:01-DQA1*01:02-DQB1*06:02* genotype maps to *DQA1*01:02-DQB1*06:02* (henceforth *DQ6*), which encodes the DQ6 molecule, rather than *DRB1*15:01-DQB1*06:02* (henceforth *DR15*)^18,19^. In our cohort, virtually everyone who carries *DQ6* also carries *DR15* (99.2% of *DQ6* positive individuals are *DR15* positive) so it is difficult to fine map the protective genotype. When both *DQ6* and *DR15* are included simultaneously in a regression analysis, *DQ6* retains significance (ln[OR]= -2.86, P=6.1x10^-3^) whereas *DR15* becomes non-significant (ln[OR]= -0.9, P= 0.4) in line with the literature^18,19^. We therefore focused our analysis on *DQ6*. However, results focusing instead on *DR15* (either as a phased haplotype or compound genotype) are virtually identical (see **SI, Supplementary Results**).

For each individual we calculated their “iKIR score”, a value equal to the count of iKIR-ligand gene pairs in that individual weighted by the strength of the iKIR-HLA interaction (**Materials and methods** and ^11^) so that a large iKIR score reflects someone with a large number of strong iKIR-ligand interactions. iKIR score is protective in our cohort (ln[OR]= -0.22, P=2.8x10^-25^). However, in a model including iKIR ligands (Bw4, C1 and C2) as covariates, the iKIR score association is lost (ln[OR]=-0.008, P=0.85) and Bw4 is strongly protective (ln[OR]=-0.45, P=9.1x10^-16^). Therefore, the iKIR score association is probably driven by KIR ligands alone; for example, by the protective or detrimental effect of *B*57:01* (Bw4-80I motif) and *A*24:02* (Bw4-80I motif) alleles respectively^20^. We conclude that iKIR score as a main effect does not contribute to T1D risk and move to investigate the impact of iKIR score on the HLA class II *DQ6* association.

On stratifying the cohort into individuals with a high iKIR score (>1.75) and individuals with a low iKIR score (≤1.75) we found that the protective effect of *DQ6* varied significantly between the strata as we had seen for class I effects in the context of virus infection, but as expected the effect was reversed with the strongest class II protection seen in individuals with a low iKIR score (**Fig. 1A**). The odds of seeing this by chance were found, by permutation test, to be P=5.6x10^-4^. The iKIR score threshold selected to categorise someone as having a “high” iKIR score (threshold=1.75) was chosen to give a balanced stratification. We investigated whether our result, that the protective effect of *DQ6* was stronger amongst people with a low iKIR score than amongst people with a high iKIR score, was dependent on the choice of threshold. The small number of cases having this protective genotype makes exploration of more extreme thresholds problematic as the number of individuals in one stratum may be quite sparse. Nevertheless, we explored 4 different thresholds: 1.5, 1.75, 2.0 and 2.5. In every case the result was replicated (**Fig. 1B**, **Table S1**). These observations are not independent, but they do demonstrate that our result is robust to the choice of threshold.

**Fig. 1.**
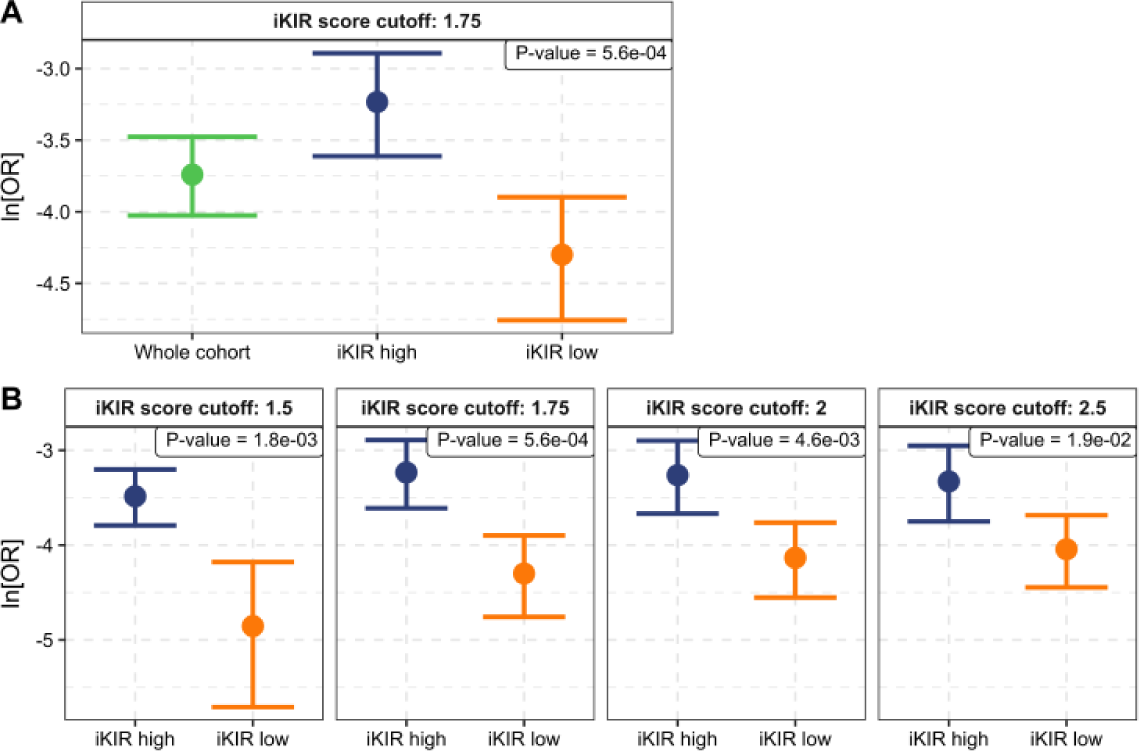
DQ6-associated protection is enhanced amongst individuals with low number of functional iKIR. **A** The protective effect of *DQ6* is enhanced in the group of individuals with an iKIR score equal to 1.75 or lower (iKIR score threshold = 1.75). The number in the top right box corresponds to the odds of seeing this enhancement by chance (Permutation test, see **Materials and methods**). **B** The same analysis was repeated using different iKIR score thresholds to define “high” and “low” (1.5, 1.75, 2.0 and 2.5, indicated in grey box above the relevant plot). The observed effect is robust to the exact choice of threshold. In all cases the dot is the natural log of the odds ratio, ln[OR], and the bars the 95% confidence intervals obtained from the logistic regression; grey: whole cohort, blue high iKIR score strata, yellow: low iKIR score strata. ln[OR]<0 indicates protection, with greater protection associated with a lower ln[OR]. Coefficients, p-values and group sizes are reported in **Table S1.**

### *DQ6* protection decreases as a function of iKIR score

If iKIR score is a meaningful measure, we might expect a “dose effect” i.e. the impact on the protective class II effect depends on the value of the iKIR score. To investigate this, rather than stratifying the cohort, we modelled the whole cohort and included iKIR score in the model as a continuous variable interacting with *DQ6 (i.e. OUTCOME* ∼ *DQ*6 × *iKIR*_*score* + *GENDER).* The interaction term was significant (P=6.57x10^-7^ model AIC= 14271); this is stronger than if we included stratified iKIR score as an interaction term (P=1.93x10^-4^ model AIC=14264) consistent with a dose effect.

To confirm that there was a dose effect we stratified the cohort into individuals with a low, intermediate, and high iKIR score and in each of the three strata calculated the protective effect of *DQ6*. Again, this analysis is problematic as the strong protective effect of *DQ6* is such that, although our cohort is very large, there are only 54 cases with the protective genotype. Due to the inevitable low numbers per strata, results would be expected to be noisy and subject to exact choice of stratification. Therefore, we considered all possible strata choices yielding 12 or more cases in each stratum. For each stratification we found the same picture, the ln[OR] decreased in a dose-dependent manner as the iKIR score decreased (**Fig. S3A**). In short, with both regression by interaction and by stratification we reached the same conclusions: *DQ6* protection decreases as a function of iKIR score.

We also assessed whether iKIR score was a better predictor of impact than iKIR count (**SI**, **Materials and methods**). The iKIR score has a marginally stronger effect than the iKIR count on the protective effect of *DQ6* (P=6.57x10^-7^ for iKIR score, P=1.1x10^-6^ for iKIR count); in backwards stepwise regression (starting from a full model with both interaction terms) iKIR count is removed from the model and the model with iKIR Count has a higher AIC than a model with iKIR score (difference=10). However, both terms are very similar and the difference in coefficient (for standardised variables) is very small (0.69 for the score 0.68 for the count) making it difficult to reach firm conclusions but on balance the effect seems to be better predicted by the iKIR score.

Additional analysis concluded that (1) the observed modification of *DQ6* cannot be explained by the HLA class I genes alone (2) all iKIR genes contribute to the *DQ6* modification (3) KIR modification of *DQ6* was most likely to be explained by inhibitory KIR rather than activating KIR (though the two are closely correlated) and (4) iKIR modification of *DQ6* is independent of the detrimental genotypes *DRB1*03:01-DQB1*02:01* and *DRB1*04:01/02/04/05-DQB1*03:02* (**Supplementary Results**).

### Other protective HLA class II *DRB1-DQB1* haplotypes are also modulated by iKIRs

Having established that a low iKIR score is associated with a significant increase in the protection conferred by the prototypical protective class II genotype *DQ6*, we hypothesized that other significantly protective haplotypes or genotypes in our cohort would also be iKIR score modified. The strongest genetic associations reported in the literature have been with *DRB1-DQB1* haplotypes, so we initially focused on phased *DRB1*-*DQB1* haplotypes.

In the whole cohort we found 17 *DRB1-DQB1* haplotypes that were significantly protective. For every case, with the exception of one haplotype (*DRB1*07:01-DQB1*03:03*), we found the same effect i.e. the protection conferred by class II haplotypes was weakened in the presence of a high iKIR score and strengthened in the presence of a low iKIR score (**Fig. S4A, Table S3**). Several haplotypes are present at low frequency in our cohort so some results may have arisen by chance. The overall probability of our observation i.e. that iKIR score modified the protection of the 17 *DRB1-DQB1* protective haplotypes was assessed by a permutation test (**Materials and methods**). The test statistic was the weighted mean of the iKIR effect (ln[OR] in KIR high – ln[OR] KIR low weighted by the haplotype frequency). We found that out of 3x10^7^ permutations there was never a case where the test statistic of the permuted dataset was as extreme as the observed value (**Fig. S4B**), so we conclude that the effect of iKIR on protective haplotypes is unlikely to have arisen by chance (odds of seeing the effect across protective haplotypes P<3x10^-7^).

Although the finding that the protective *DRB1-DQB1* haplotypes were significantly modified by functional iKIRs is interesting we were aware of two potential caveats. First, the *DR-DQ* haplotypes may not be the causal drivers of protection, they could just be neutral passengers marking class II genotypes that are more closely associated with protection (i.e. we are focusing on the wrong target). Second, linkage disequilibrium between the class II haplotypes and protective or detrimental HLA class I alleles (which are also iKIR ligands) could be mistaken for iKIR modification of the class II protective effect. We therefore investigated both these possibilities.

### Which alleles are best associated with protection?

We suspected that some of the *DRB1-DQB1* haplotypes may not be protective themselves but that they were neutral passengers marking the true, causal driver genotypes. Therefore, we aimed to understand whether the *DRB1* and *DQB1* genes of the protective haplotype themselves or other class I or class II alleles best marked the protective effect (and if the latter whether these protective alleles were also modulated by iKIR score).

We investigated all HLA class I and HLA class II alleles as well as two and three allele genotypes at *DRB1*, *DQA1* and *DQB1* (considered in cis and in trans as both trans-acting and cis-acting associations have been documented^4^). We considered all pairs of genotypes in a regression model (a total of 235,347,360 pairwise combinations) and defined “drivers” to be genotypes that never lost significance nor swapped direction in the presence of another genotype. Of course, we cannot rule out the possibility that these genotypes mark unsequenced variants that are even more significantly associated with outcome, but we can say they are the most significantly associated of the class I and class II alleles. We identified 10 HLA class I alleles and 21 class II genotypes (single alleles, pairs or trios) in our cohort significantly associated with T1D independently of all other genotypes in the cohort (**Fig. 2**Fig. 2. Forest plot with all HLA drivers associated with T1D., **Table S4**). Establishing a list of the class II driver genotypes is difficult due to the strong linkage disequilibrium across the HLA region and in the case of colinear or close to colinear genotypes, simplifying assumptions had to be made (see ***SI, Materials and methods***). We adjusted for multiple comparisons using the effective number of tests (M_eff_=3,692, calculated from the correlation structure of the original 21,696 genotypes considered, ***SI, Materials and methods***) and applied the Bonferroni correction (0.05/M_eff_); this gave a cutoff for significance of P<1.35x10^-5^. All 31 driver genotypes remained significant. This remained true even when assuming that all genotypes tested are independent (m=21,696, threshold P=0.05/m=2.3x10^-6^) or when using the typical significance level in GWAS studies (P< 5x10^-8^).

**Fig. 2.**
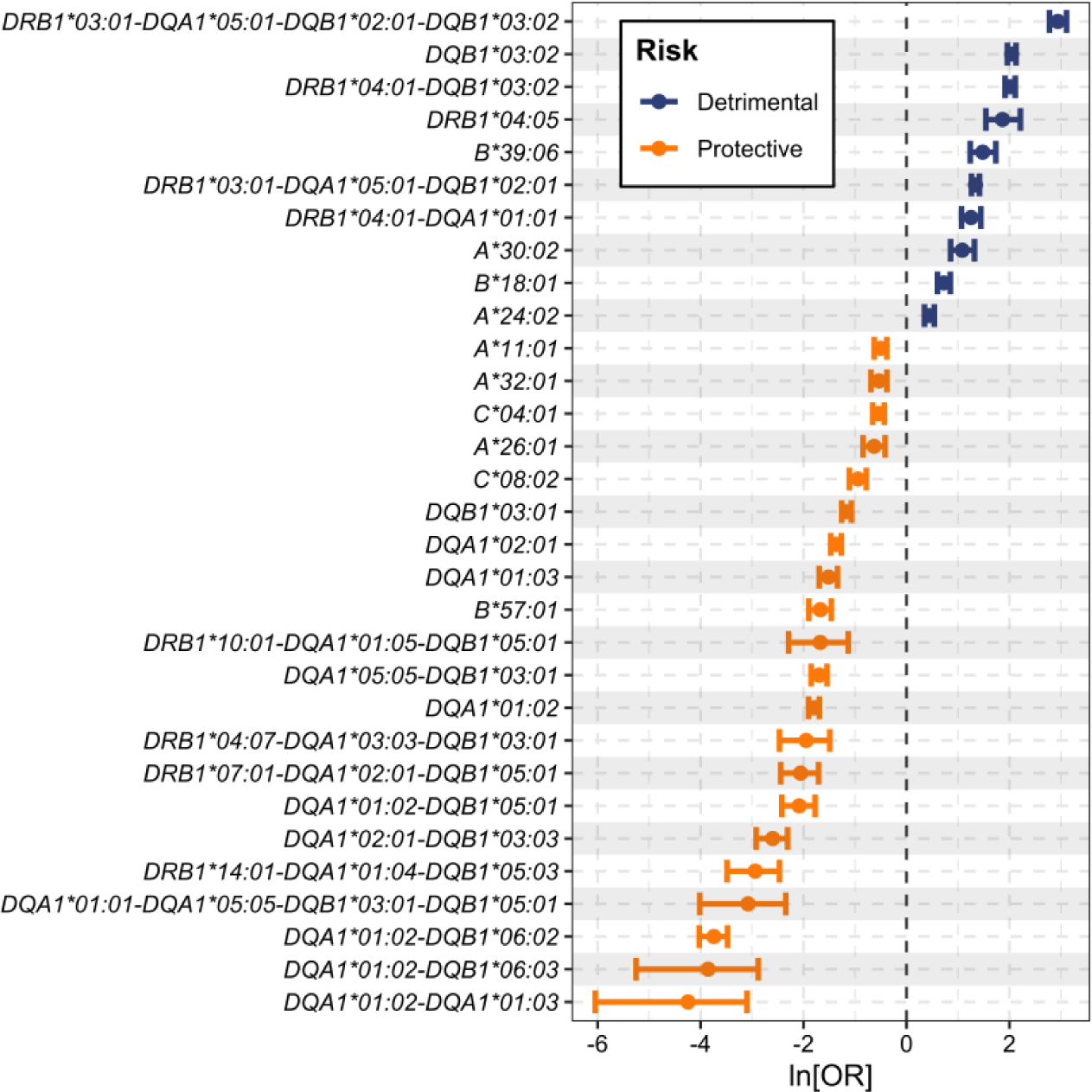
Forest plot with all HLA drivers associated with T1D. We show all class I and class II driver genotypes identified (see ***SI, Materials and methods***). The regression coefficients plotted, i.e. the ln[OR], are for an analysis in the GRID cohort in which the only other covariate was gender. The coefficients will depend both on the genetic risk of the whole cohort (i.e. the background that the risk/protection is measured relative to) and the other genotypes correlated with the genotype of interest. Note that although *DRB1*03:01-DQA1*05:01-DQB1*02:01-DQB1*03:02* appears to be a compound of the detrimental genotype *DRB1*03:01-DQA1*05:01-DQB1*02:01* and the detrimental genotype *DQB1*03:02* it was retained in the list of independent drivers as it retained direction and significance of effect (albeit considerably weakened) in multiple regression when both *DRB1*03:01-DQA1*05:01-DQB1*02:01* and *DQB1*03:02* were included simultaneously with it. We cannot rule out the possibility that these genotypes mark unsequenced variants that are even more closely identified with outcome, but we can say they are the class I and class II alleles most closely associated with outcome. Coefficients, p-values and number of cases and controls are provided in **Table S3**.

As anticipated, none of the 17 protective *DRB1-DQB1* haplotypes studied above were the drivers of protection. Instead, we identified 15 protective HLA class II genotypes which were better associated with outcome. Henceforth we focus on these 15 protective genotypes.

### Impact of HLA class I drivers

Our second concern was that correlations between the protective class II genotypes of interest and the driver class I genotypes (some of which are also iKIR ligands) could be mistaken for iKIR modification. For example, *HLA-B*57:01* is a class I allele which is significantly associated with protection. It also encodes the ligand for KIR3DL1 and as such will be enriched in the iKIR-high strata. If a protective class II genotype is associated with *B*57:01* then individuals with the class II genotype and *B*57:01* will be more likely to appear in the high iKIR strata; any additive effects of protection from the class II genotype and *B*57:01* would then risk either being misinterpreted as iKIR modification of the class II genotype or risk masking an iKIR modification (depending on the direction of the correlations). To remove this possibility, we removed all individuals who were positive for any of the class I driver alleles from the cohort, leaving a reduced cohort of size N=5,420. Simply removing class I drivers is the cleanest approach to dealing with the problem. In many immunogenetics analyses this is not an option due to the reduction in cohort size; in this case we are fortunate in starting with a very large cohort which allows removal of the class I drivers.

### Protective effects of HLA class II genotypes are significantly modified by iKIR

We stratified this reduced cohort into individuals with high iKIR score and individuals with a low iKIR score and analyzed the effect of the protective class II drivers within each stratum; class II drivers which were carried by fewer than 10 cases were not studied as numbers were too low to permit stratification. The nine remaining protective genotypes were all modified in the same direction at all three thresholds considered (**Fig. 3A**, **Table 1**). The only genotypes for which the iKIR effect was not extremely clear (*DQA1*0201-DQB1*03:03* and *DQA1*01:02-DQB1*05:01*) were very infrequent (N=12 and N=14 cases respectively) so the number of carriers in each strata would be very low possibly explaining the lack of a clear modification. Overall, the odds of observing this iKIR modification across the 9 genotypes was very low, P=2x10^-8^ (**Fig. 3B**). Furthermore, we also modelled the whole cohort and included iKIR score as an interaction term with a given protective genotype. The iKIR interaction term was significant in the 4 out of 5 most frequent genotypes and in the expected direction in all genotypes (**Table S5**). We conclude that iKIR negatively impacts the effects of all protective class II genotypes in T1D, that this effect is attributable to iKIR-HLA receptor ligand interactions and unlikely to be observed by chance.

**Fig. 3.**
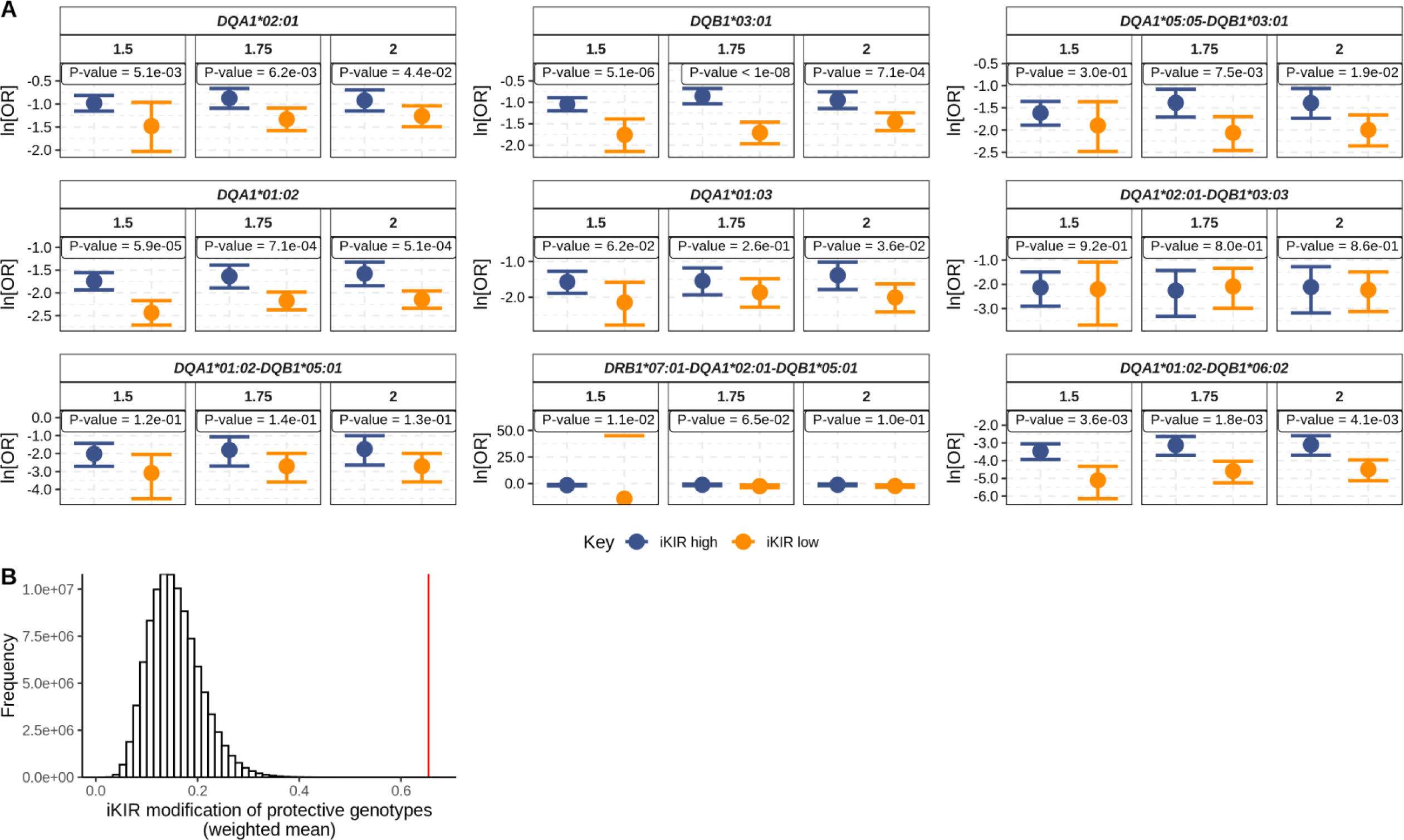
iKIR score negatively impacts the protective effect of 9 driver HLA class II genotypes in T1D. **A** The 9 protective class II genotypes investigated were all significantly modified by iKIR at the three thresholds (definition of iKIR high and low) considered. The only two genotypes for which the effect was not strong (*DQA1*0201-DQB1*03:03* and *DQA1*01:02-DQB1*05:01*) were very infrequent (N=12 and N=14 cases respectively). **B** The observed value of our test statistic (weighted mean of the difference in ln[OR] between the KIR high and the KIR low strata at threshold=1.75), indicated by the red line, lies far above the distribution (grey histogram) of the same test statistic under the null hypothesis that the iKIR score has no impact on the protective genotypes (generated by permuting the iKIR score of individuals in the cohort). Indicating that the probability of obtaining our observation by chance is extremely low (P=2x10^-8^).

**Table 1.** iKIR score decreases protection associated with protective class II genotypes in T1D. iKIR score decreases protection associated with protective class II genotypes in T1D. The GRID cohort without carriers of HLA class I drivers (N=5,420) was stratified into individuals with high or low iKIR score at a 1.75 cutoff (we also tested cutoffs 1.5 and 2.0, see Table S6). The protective effect of each genotype was evaluated independently in each stratum using multivariate logistic regression with gender as covariate. HLA class II protection is enhanced in the iKIR low strata for all genotypes but for the very infrequent protective genotype *DQA1*02:01-DQB1*03:03*. Regression coefficients, permutation p-values and cohort sizes are reported for the different strata. P-value for the whole cohort (unstratified analysis) calculated using the Wald-test; p-values for the stratification analysis are calculated using the permutation test.

| Genotype | Group | ln[OR] | 2.50% | 97.50% | P-value | N Genotype + |  | N Genotype - |  |
| --- | --- | --- | --- | --- | --- | --- | --- | --- | --- |
|  |  |  |  |  |  | Cases | Controls | Cases | Controls |
| **DQA1\*01:02-DQB1\*06:02** | Whole cohort | -3.91 | -4.33 | -3.53 | 1.76x10<sup>-83</sup> | 26 | 729 | 2984 | 1681 |
|  | iKIR high | -3.12 | -3.69 | -2.63 | 1.8x10<sup>-3</sup> | 15 | 279 | 1072 | 882 |
|  | iKIR low | -4.58 | -5.25 | -4.03 |  | 11 | 450 | 1912 | 799 |
| **DQA1\*02:01-DQB1\*03:03** | Whole cohort | -2.21 | -2.87 | -1.65 | 9.05x10<sup>-13</sup> | 12 | 85 | 2998 | 2325 |
|  | iKIR high | -2.25 | -3.31 | -1.43 | 0.8 | 5 | 49 | 1082 | 1112 |
|  | iKIR low | -2.09 | -2.99 | -1.34 |  | 7 | 36 | 1916 | 1213 |
| **DQA1\*01:02-DQB1\*05:01** | Whole cohort | -2.32 | -2.92 | -1.80 | 4.14x10<sup>-16</sup> | 14 | 109 | 2996 | 2301 |
|  | iKIR high | -1.80 | -2.69 | -1.06 | 0.14 | 7 | 44 | 1080 | 1117 |
|  | iKIR low | -2.71 | -3.58 | -1.99 |  | 7 | 65 | 1916 | 1184 |
| **DRB1\*07:01-DQA1\*02:01-DQB1\*05:01** | Whole cohort | -1.69 | -2.37 | -1.10 | 1.40x10<sup>-7</sup> | 12 | 52 | 2998 | 2358 |
|  | iKIR high | -1.13 | -1.94 | -0.41 | 0.065 | 9 | 29 | 1078 | 1132 |
|  | iKIR low | -2.45 | -3.89 | -1.39 |  | 3 | 23 | 1920 | 1226 |
| **DQA1\*01:02** | Whole cohort | -1.91 | -2.07 | -1.76 | 8.14x10<sup>-133</sup> | 252 | 922 | 2758 | 1488 |
|  | iKIR high | -1.63 | -1.88 | -1.38 | 7.1x10<sup>-4</sup> | 88 | 359 | 999 | 802 |
|  | iKIR low | -2.18 | -2.38 | -1.98 |  | 164 | 563 | 1759 | 686 |
| **DQA1\*05:05-DQB1\*03:01** | Whole cohort | -1.74 | -1.99 | -1.51 | 6.015x10<sup>-46</sup> | 88 | 355 | 2922 | 2055 |
|  | iKIR high | -1.39 | -1.72 | -1.08 | 7.5x10<sup>-3</sup> | 53 | 198 | 1034 | 963 |
|  | iKIR low | -2.05 | -2.44 | -1.69 |  | 35 | 157 | 1888 | 1092 |
| **DQA1\*01:03** | Whole cohort | -1.75 | -2.03 | -1.49 | 2.12x10<sup>-36</sup> | 67 | 279 | 2943 | 2131 |
|  | iKIR high | -1.54 | -1.93 | -1.18 | 0.25 | 35 | 156 | 1052 | 1005 |
|  | iKIR low | -1.86 | -2.27 | -1.48 |  | 32 | 123 | 1891 | 1126 |
| **DQA1\*02:01** | Whole cohort | -1.15 | -1.31 | -0.99 | 2.02x10<sup>-45</sup> | 256 | 548 | 2754 | 1862 |
|  | iKIR high | -0.88 | -1.10 | -0.67 | 6.2x10<sup>-3</sup> | 149 | 322 | 938 | 839 |
|  | iKIR low | -1.32 | -1.57 | -1.08 |  | 107 | 226 | 1816 | 1023 |
| **DQB1\*03:01** | Whole cohort | -1.25 | -1.39 | -1.11 | 5.59x10<sup>-69</sup> | 361 | 777 | 2649 | 1633 |
|  | iKIR high | -0.85 | -1.03 | -0.67 | <1x10<sup>-8</sup> | 269 | 505 | 818 | 656 |
|  | iKIR low | -1.71 | -1.96 | -1.46 |  | 92 | 272 | 1831 | 977 |

### iKIR score modification is replicated in an independent cohort

To validate our findings, we studied an independent dataset consisting of 339 US multiplex families from the HBDI consortium. Prior to analyzing this cohort, we conducted a power analysis (***SI, Materials and methods***) to assess whether we had sufficient statistical power to detect a significant iKIR score effect in this much smaller cohort. We found that, assuming the effect size was the same as in the GRID cohort, the family cohort would not be sufficiently powered to detect a significant iKIR score modification of individual protective class II genotype (**Fig. S5A**). However, we estimated that there was sufficient power to detect an iKIR score modification if we considered several protective genotypes simultaneously i.e. *DQB1*03:01*, *DQA1*02:01*, *DQA1*01:02* and *DQA1*01:02-DQB1*06:02* (**Fig. S5B**) and analyzed the family dataset on this basis. We stratified the family cohort into trios with a high iKIR score (threshold>1.75) and trios with a low iKIR score (threshold <1.75) based on the iKIR score of the affected child. For each genotype, we calculated the difference between the ratio of transmissions and non-transmissions in each strata (see ***SI, Materials and methods***). The odds of observing an equal or greater difference between the ratios in each stratum under the null hypothesis was statistically significant P=1x10^-5^ (Permutation test). These conclusions remained unchanged when using different iKIR score thresholds (**Table S7**). We conclude that our finding that iKIR score impacts HLA class II mediated protection was replicated in an independent dataset.

### Fraction of cases prevented by a low number of functional iKIR

To quantify the impact of functional iKIR on disease prevalence we estimated the fraction of cases of T1D prevented by the iKIR interaction with the most protective genotype (*DQ6*) using data from the GRID cohort and the prevalence of T1D in Europe^21^. If the population all carried a high number of functional iKIR (i.e. iKIR score>1.75) then we estimate that the cases prevented by *DQ6* would be 21.6% but if the population all had a low number of functional iKIR (i.e. iKIR score ≤ 1.75) then 31.6% of cases would be prevented, an increase of more than 45%. To put this iKIR effect in context of other T1D-associated SNPs we normalized the iKIR score so it was on a scale of [0,1] (in line with presence/absence of a variant SNP) and then considered it as a main effect in a *DQ6+* cohort. The OR for iKIR score was 6.12 with a 95% confidence interval (CI) of 1.96 to 19.69 (and OR=7.55, 95%CI 1.61-37.21 in the smaller cohort with individuals carrying HLA class I drivers removed). The more conservative value is plotted alongside the OR of other variants which have previously been associated with T1D for comparison (**Fig. 4**).

**Fig. 4.**
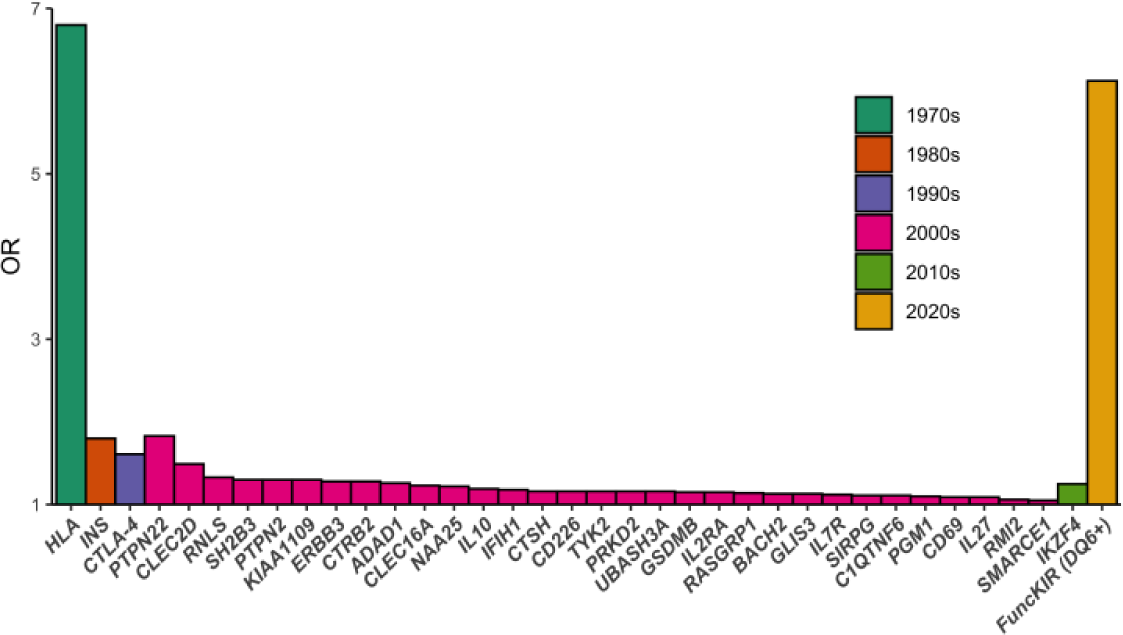
Loci that affect the risk of T1D. The OR for functional iKIR score in a *DQ6*+ cohort is shown (orange) alongside the OR for variants in other loci as reported in the literature (***SI, Materials and Methods***). Associations are grouped (and colour-coded) by decade when the association was first reported (not necessarily fine-mapped) and then within each decade, associations are ranked by size of OR. Gene names refer to most likely candidate in the region. Figure is after a similar figure in Rich *et. al.*^58^.

### Mathematical model of the β-cell autoimmune destruction can recapitulate the iKIR score effect in the presence of saturation

If iKIRs increase CD4+ T cell lifespan (either directly via iKIR expression on the affected T cell or indirectly via iKIR expression on NK cells), as we have reported for CD8+ T cells^8^, one might hypothesize that an iKIR-mediated enhancement of autoreactive T cell survival is detrimental, which agrees with our immunogenetic findings on protective HLA genotypes. However, by this reasoning, one would expect that the risk conferred by detrimental HLA genotypes like *DRB1*04:01-DQB1*03:02* would also be modulated by iKIR score, i.e., higher risk in individuals carrying detrimental HLA genotypes and high number of iKIR genes. To investigate this apparent discrepancy in the immunogenetic results we use mathematical modelling. Briefly, we implemented an ordinary differential equation system that reflects the interactions between the T cells and insulin producing β-cells in the pancreatic islet based on an existing model of the human antitumor T cell response^22^. We implemented two versions of this model, with and without density-dependent T cell production (see ***SI, Materials and methods***) and generated a cohort of 10,000 in silico individuals, each one carrying a parameter combination randomly sampled from parameter ranges obtained from the literature (**Table S15**). For each of the two models and for each parameter set, we run the simulation twice to mimic the iKIR positive effect on T cell survival^11,23^; the first simulation has lower T cell death rates (iKIR high) than the second one (iKIR low). In both models, increasing T cell survival resulted in progression to T1D in a small fraction of in silico individuals who would otherwise have been healthy but for the majority of simulations the outcome (health or development of T1D) was independent of iKIR (**Fig. 5A-B**). We then asked whether an increased T cell survival is associated with higher T1D risk in carriers of protective but not neutral nor detrimental HLA genotypes. We assumed that HLA class II-protected in silico individuals have high number of islet-specific Tregs, as reported recently in a study on healthy individuals^24^. We classified individuals into groups on the basis of mean Treg numbers during the simulation and then computed the difference in ln[OR]s between the iKIR high and iKIR low in silico cohorts in each group. In model 1 (without density dependent T cell production), the difference of lnORs remained constant for different groups with different numbers of Tregs (**Fig. 5C**). In model 2 (with density dependent T cell production) though, the difference in ln[OR]s increases as the number of Tregs per islet increases (**Fig. 5D**), which recapitulates the trend observed in the actual data (**Fig. 5E)** and provides a possible explanation to our seemingly contradictory immunogenetic results. When Treg levels are saturated and reach carrying capacity – i.e., in protective HLA carriers – the increase in T cell survival results in an increase of Tconv but not Treg population size. Consequently, in this scenario, there is an effective increase of β-cell destruction that cannot be compensated by Treg suppression of Tconvs. In unsaturated conditions though – i.e., in neutral or detrimental HLA carriers – iKIR-mediated increase of both Treg and Tconv survival results in a zero net effect on β-cell destruction. These results are consistent with our observations.

**Fig. 5.**
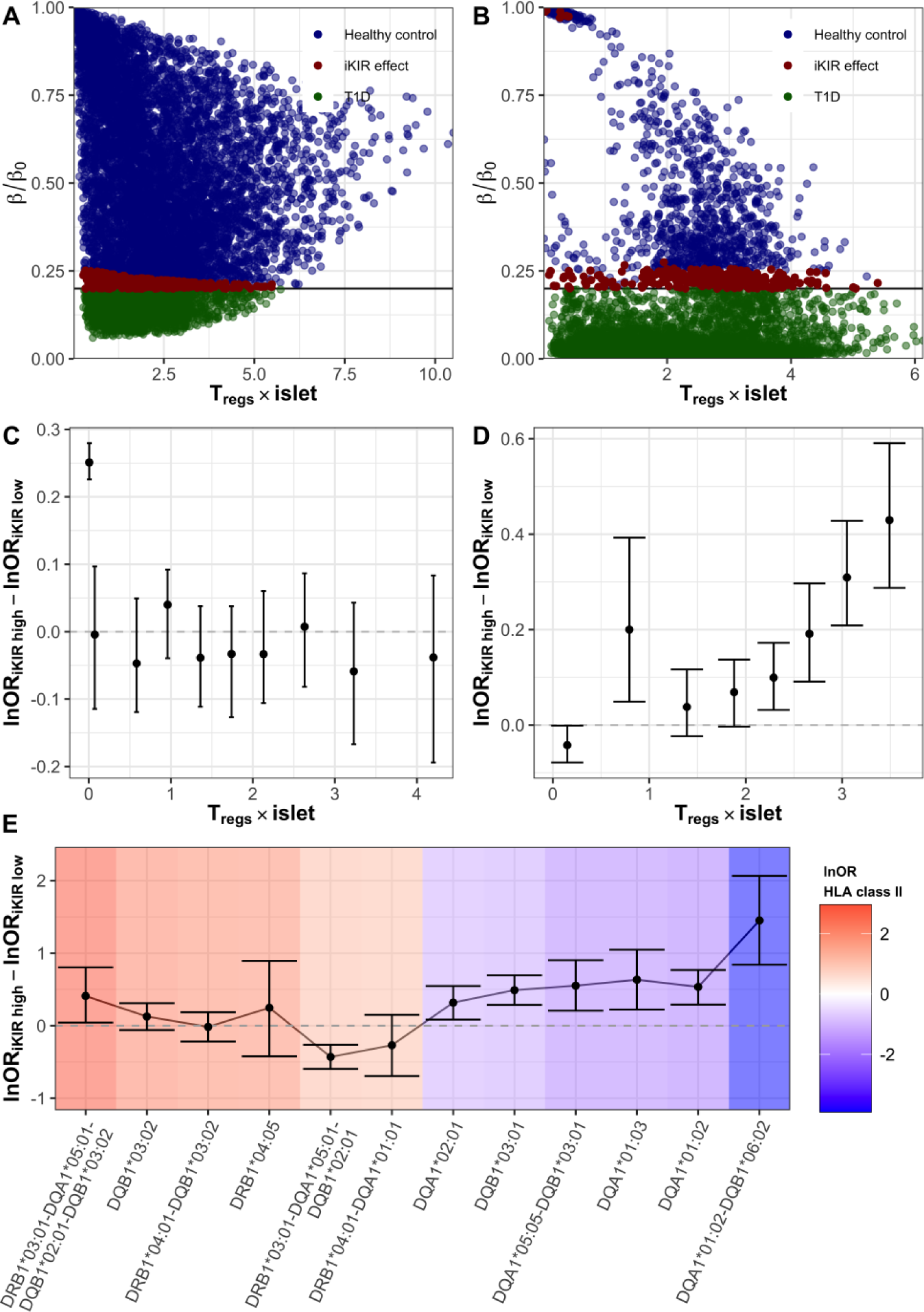
Mathematical model. Simulations of iKIR effect on the T cell response against insulin-producing beta cells. **A** and **B** Fraction of beta cell mass remaining as a function of the number of regulatory Tregs during the immune response is shown for each in silico individual in the simulated cohort with model 1 (**A**) and model 2 (**B**). Color code indicates outcome of the immune response: individuals that remain healthy after the immune response are shown in blue, those that transition to T1D are shown in green and individuals that have a different outcome depending on their functional iKIR gene count are shown in red. Note that the outcome depends on the threshold of disease onset. **C** and **D** Difference of OR for a cohort with high number vs a cohort with low number of functional iKIR genes as a function of number of Tregs during the immune response simulated with model 1 (**C**) and model 2 (**D**). E Difference between the ln[OR] of HLA class II genotypes in the group of GRID individuals with a high iKIR score and the group with a low iKIR score. Color code according to the lnOR of the HLA class II genotype (risk).

### KIR+ T cell frequency is not increased in T1D patients

There is experimental evidence showing that iKIRs impact T cell responses via two main pathways. Directly, the ligation of iKIRs enhances T cell survival in vitro^9,11^. Indirectly, NK cells also modulate T cell lifespan by regulating activated T cell numbers. Recent work supports the latter pathway as being the most relevant in healthy and virus-infected individuals^8^. We wanted to investigate whether this was also the case in T1D. We hypothesized that if the functional iKIR effect on HLA class II genotypes is caused by the expression of iKIRs on T cells (direct pathway), we would expect to see differential expression of iKIRs on T cells between T1D patients and healthy controls. To test this hypothesis, we analysed scRNAseq data from PBMCs samples of 4 children with islet auto-antibodies (two of them developed T1D by 36 months of age) and 4 matched controls (see **Materials and methods**). Only two barcodes labelled as terminal effector CD4+ T cells were positive for *KIR* transcripts and none of the cells labelled as Tregs expressed KIRs. As expected, a greater proportion of terminally differentiated CD8+ T cells expressed KIRs (**Fig. 6A**). Nevertheless, we detected *KIR* transcripts in CD8+ T cells from both seropositive individuals and healthy controls, suggesting that KIR expression in blood is not altered in disease. To validate those findings, we recruited 10 T1D patients (including new onset T1D patients and individuals with long standing disease) and 10 matched healthy controls and performed KIR immunophenotyping of CD4+, CD8+ and NK cell subsets by flow cytometry (see **Materials and methods**). We found that KIR protein expression was higher in late stage differentiated T cells and for KIR2DL2/L3 compared to KIR2DL1, in agreement previous findings^8^. As in the scRNAseq analysis, we did not observe differences in KIR expression between T1D cases and controls within naïve or memory subsets (**Fig. 6B**). As expected, KIR+ CD4+ T cells were rare, and frequencies were again comparable between cases and controls (**Fig. 6C**). In summary, the frequency of KIR+ T cells is not altered through T1D disease stages, which argues against a direct effect of KIRs on T cells as the underlying mechanism of the iKIR gene modulation on HLA associations, consistent with our previous findings. We suggest, in line with the evidence for CD8+ T cells^8^, that iKIRs enhance CD4+ T cell survival via the indirect pathway.

**Fig. 6.**
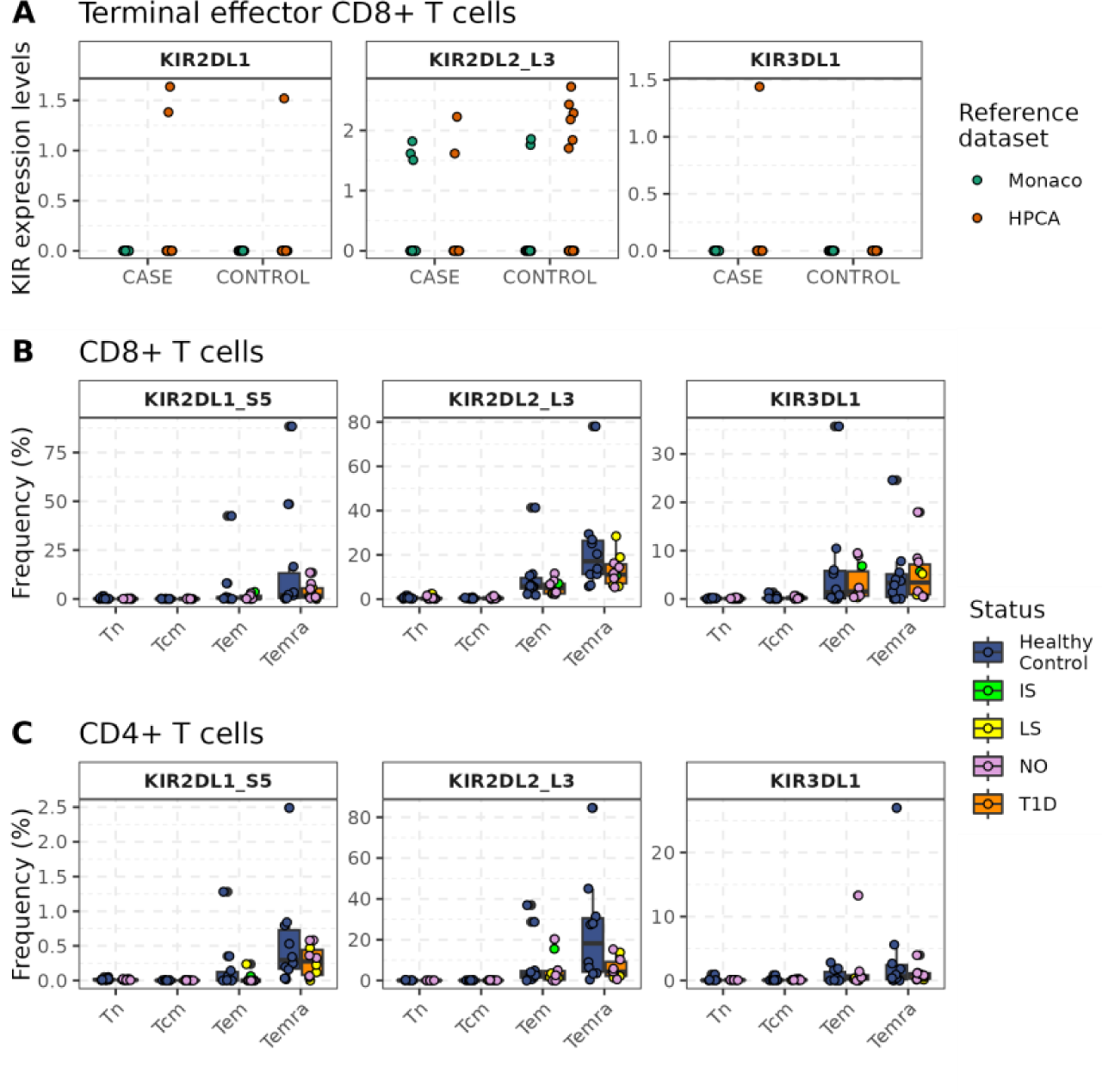
Frequency of terminal effector CD8+ T expressing KIR transcripts is similar between seropositive children and healthy controls. **A** Few CD8+ terminal effector memory cells express KIR transcripts. Cells labelled as *Terminal effector CD8 T cells* (Monaco reference, green) or as *T cell:CD8+ effector memory RA* (HPCA reference, orange) are shown split by disease status (case, control) and by KIR gene (*KIR2DL1*, *KIR2DL2/L3* and *KIR3DL1*). **B** The percentage of cells in each CD8+ T cell population (Tnaive, Tem, Tcm, Temra) expressing iKIRs quantified by flow cytometry. **C** The percentage of cells in each CD4+ T cell population (Tnaive, Tem, Tcm, Temra) expressing iKIRs quantified by flow cytometry. **B, C** Dots represent frequencies for each individual in the cohort. T1D samples are colour coded according to disease duration at time of collection (NO=new onset, IS=intermediate standing disease, LS=long standing disease). Boxes show medians and interquartile ranges within T1D individuals (N=10, orange, irrespective of disease duration) and healthy individuals (N=10, blue).

## Discussion

Our aim was to conduct an immunogenetic analysis to study the role of iKIRs in autoimmunity. Specifically, we wanted to investigate whether functional iKIR genes have a significant impact on HLA associations in T1D. This interaction is clinically significant in other contexts; we previously reported that functional iKIR genes enhanced protective and detrimental HLA disease associations in chronic viral infections. We postulated that a similar effect might be relevant in autoimmunity. We have analyzed a large (N=11,961) case-control T1D dataset to investigate the effect of functional iKIR genes on HLA class II genetic associations. We found that a low number of functional iKIR genes (iKIR genes together with the genes encoding their corresponding HLA class I ligands), enhanced the dominant protection conferred by *DQA1*01:02-DQB1*06:02*. The effect was driven by iKIR-ligand pairs rather than HLA ligands alone, which are independently associated with T1D. The same results were observed for the other 9 protective HLA class II genotypes in our cohort; for all but one infrequent genotype, the results were statistically significant and observed at all iKIR stratifications considered. The odds of observing this effect across all protective genotypes by chance is low (P=2x10^-8^). Moving onto an independent replication cohort, an identical result was observed with protective class II genotypes being more protective in individuals with a low number of functional iKIR genes. Again, the probability of observing this result by chance was low (P=1x10^-5^). In striking contrast, functional iKIR had no consistent impact on detrimental class II associations. Most detrimental genotypes showed no iKIR modification at all. Two detrimental genotypes which were iKIR modified were modified in opposite directions and on closer examination this was found to be attributable to linkage disequilibrium with a class I genotype in one case and inverse correlation with a protective class II genotype in the other case. In short, whilst the functional iKIR modification of protective HLA class II associations is very clear and highly statistically significant (UK-GRID cohort: P=2x10^-8^, replication cohort: P=1x10^-5^), the absence of an iKIR modification of detrimental genotypes is equally clear (P=0.46).

We studied the underlying mechanism using single cell RNAseq data, protein expression data and mathematical modelling. Evidence from ours and others studies indicate that iKIR ligation increases T cell lifespan. Recently we have shown that, for CD8+ T cells, iKIR expression by a third cell (other than the CD8+ T cell whose lifespan is extended) is necessary and that iKIR expression by the CD8+ T cell of interest is not relevant. If iKIR expression on T cells did explain our observations in T1D then iKIR gene expression on T cells might be expected to differ between T1D patients and healthy individuals. We did not find major differences in the size of KIR+ immune populations between cases and controls, suggesting, as hypothesized, that the direct ligation of iKIRs on T cells is not the main underlying mechanism of the functional iKIR gene effect. This is consistent with our recent work which also effectively rules out direction expression of iKIRs on CD8+ T cells as a mechanism for those T cells increased survival^8^. Finally, we used a simple mathematical model of the immune destruction of β-cells to generate plausible hypotheses about the iKIR modulation of protective but not neutral nor detrimental HLA associations. We predicted that an increase of T cell survival – driven by a high number of functional iKIR genes – would have a detrimental effect only when regulatory T cells are present at saturation levels, corresponding to individuals with a protective HLA class II genotype. When T cell numbers are far from the T cell population carrying capacity, a longer T cell lifespan has a zero net effect on beta cell destruction; a change in conventional T cell numbers is compensated by a change in regulatory T cell numbers.

Although HLA class II associations with T1D are very well studied, to the best of our knowledge there are no reports of how these associations are modified by functional iKIR genes. There are several studies investigating iKIR genes and/or functional iKIR genes in T1D^13-15,25-27^ and a very large number of studies investigating HLA class II associations (e.g. ^2,4,28^) but none reporting the interaction. Although the absence of evidence for an interaction between functional iKIR genes and HLA class II genes in candidate gene studies can easily be explained by the argument that no one was motivated to study this particular three-way genetic association, it might be wondered why it was not picked up in one of the very large “catch-all” genome wide association studies (GWAS) performed in T1D^28-30^. The reason is that three gene associations of the type we report (KIR-ligand-class II) are never studied in GWAS as the explosion in the number of multiple comparisons for comparing all triplets of variants is prohibitive and so it is to be expected that the functional iKIR modification of HLA class II associations which we report would not be found by GWAS. Whilst we are not aware of studies of functional iKIR modifications of HLA associations in T1D, we have previously reported functional iKIR modification of protective and detrimental HLA class I associations in three chronic viral infections: human immunodeficiency virus type 1 (HIV-1), hepatitis C virus and human T cell leukemia virus type 1 (HTLV-1)^11,12^.

There are interesting parallels between these previous studies in chronic viral infection and this current study in T1D in that all show highly significant functional iKIR modification of protective HLA associations that cannot be explained just by KIR or just by class I ligands. However, in both HIV-1 and HTLV-1 infection, detrimental HLA class I genotypes (*HLA-B*35* and *HLA-B*54* respectively) were significantly modified^31^ whereas in T1D there was no evidence for functional iKIR modification of detrimental genotypes. Using mathematical modelling, we found a possible explanation for the preferential iKIR modification of protective HLA associations in T1D. If we assume that the most detrimental compound genotype *DRB1*03:01-DQA1*05:01-DQB1*02:01-DQB1*03:02* fails to produce the necessary islet-specific Tregs whereas the dominant protective genotype *DQA1*01:02-DQB1*06:02* drives Treg levels close to carrying capacity, then the iKIR effect is only manifest in the highly protective end of the spectrum of HLA class II associations. In summary, even when assuming that protective and detrimental HLA class II associations in T1D affect a common pathway, Treg numbers, we show that there is a possible mechanism that can recapitulate our observations in the data.

Broadly speaking there are two (non-exclusive) ways in which iKIR could affect class II-restricted CD4+ T cells: either by iKIR expression directly on CD4+ T cells or indirectly by iKIR expression on another population (that interacts with APCs expressing class II or CD4+ T cells restricted by class II). Here we discuss these two possibilities in turn starting with the “direct” hypothesis. Ligation of iKIRs expressed on the surface of T cells leads to phosphorylation of ITIMs in their cytoplasmic tail which recruits phosphatases including SHP1 leading to inhibition of TCR signalling which in turn can decrease effector function including cytokine production^32^ and regulation^33^ or modulate differentiation^34^; iKIRs on T cells have also been shown in vitro and in murine models to prolong CD8+ and CD4+ T cell lifetime^9,11,23,35,36^. Qin et al have reported that KIR3DL1 expression on Tregs negatively regulates Treg function in the NOD mouse and promotes T1D^33^ which could be a plausible mechanism underlying our immunogenetic findings. Furthermore, in ankylosing spondylitis, cumulative evidence indicates ligation of KIR3DL2 on Th17 CD4+ T cells may promote their accumulation and survival^10,37,38^. Finally, there are interesting parallels with other inhibitors of T cell signalling including lymphoid protein tyrosine phosphatase and PD-1. Lymphoid protein tyrosine phosphatase (LYP) is a phosphatase which, like the iKIR, negatively regulates T cell receptor signalling. A SNP (1858 C->T) in *PTPN22,* which encodes LYP, is significantly associated with T1D. The disease associated variant is associated with stronger T cell inhibition^39,40^ and Tregs from donors homozygous for the variant have decreased ability to regulate other T cells compared to Tregs from a donor homozygous for the major allele^41^. Analogously, we find greatest disease risk amongst individuals with a high number of functional iKIR genes. Similarly, PD-1 is an inhibitory receptor expressed by T cells with a similar downstream signalling pathway to iKIR, and is known to play a role in peripheral tolerance and regulation of autoimmunity^42^. Perhaps the strongest argument against a direct effect of iKIRs on CD4+ T cells is the extremely low numbers of CD4+ T cells expressing iKIRs. In healthy homeostasis only about 0.1-1% of memory CD4+ T cells express iKIR. It is hard to see how such a small population could have such a large biological effect. Furthermore, we found that iKIR expression on CD4+ cells was not increased in patients with T1D. This finding is in contrast with other autoimmune diseases. For example, iKIR expression by CD4+ T cells is increased in ankylosing spondylitis (1-6% of all CD4+ cells are KIR3DL2+ , rising to 10-60% of Th17 CD4+ cells^10^) and other autoimmune diseases such as lupus show similar increases in iKIR expression^43^. A more recent study reports increased KIR+ T cell frequencies not only in lupus patients but also in individuals with multiple sclerosis and coeliac disease^44^. In T1D, we found that both KIR gene expression and protein expression is rather similar between cases and controls. This is true for both individuals with long standing and recent onset disease as well as for individuals without clinical diagnosis but positive for islet auto-antibodies.

The alternative, “indirect” hypothesis posits that iKIRs on another cell population indirectly modulate CD4+ T cells restricted by protective class II molecules. Probably the most likely contender for such a population is NK cells as they express high levels of iKIR and are known to interact with APCs and T cells both of which could lead to a modulation of class II restricted CD4+ cell responses. NK cells kill autologous, activated but not resting CD4+ and CD8+ T cells^45,46^. *In vitro* experiments show that activated CD4+ T cells upregulate HLA-E, the ligand for the inhibitory receptor CD94-NKG2A to protect themselves from NK cell killing^47,48^. The same inhibitory receptor plays a role in the experimental autoimmune encephalomyelitis (EAE) mouse model for multiple sclerosis^49^. In this model, prevention of engagement of CD94-NKG2A, either by antibody-blockade or by knockin of a mutated ligand, resulted in elimination of autoreactive T cells and improvement of EAE. Furthermore, in studies of a murine T1D model, NOD mice immunized with Complete Freund’s adjuvant, NK cells decreased the numbers of autoreactive CD8+ T cells preventing T1D^50^. Similarly, NK cells can suppress CD8+ T cell-mediated hyperglycemia in a mouse model characterized by the transgenic expression of lymphocytic choriomeningitis virus on β-cells^51^. Perhaps, NK cells and Treg cells, act in synergy to suppress autoreactive T cell responses and prevent autoimmunity. How functional iKIR modify HLA associations in T1D and whether similar modifications are seen in other autoimmune disease are exciting and important areas that merit further research.

One limitation of this work is that the iKIR-ligand binding groups which we use in our definition of “functional iKIR” are simplistic. Incomplete knowledge of how different KIR and HLA alleles and different (HLA bound) peptides affect KIR-HLA binding and signalling precludes a more sophisticated definition. Nevertheless, these simple groupings have proved very powerful in other studies^52-57^. With this definition of inhibitory score, we saw clear and reproducible results both in the T1D cohorts and in the HIV-1, HCV and HTLV-1 cohorts. This suggests that the inhibitory score is a meaningful metric. It is worth noting that in the majority of the analyses, the score is only used to split the cohorts in half, so second order changes to the calculation of the score will not necessarily change the results (because a person’s score can change considerably, and they will still remain in the same strata).

The importance of this work is twofold. First, we have identified a family of genes that significantly impact T1D risk. We estimate that, even if we only consider a single protective class II genotype, a population with a low number of functional iKIR would see more than a 45% increase in the fraction of cases prevented compared to a population with a high iKIR score. This constitutes one of the largest genetic risk factors for T1D reported in recent decades. Second, we find evidence that iKIRs have an impact on the T cell response in vivo. A number of in vitro and murine studies indicate that iKIRs can modulate T cell responses in autoimmunity; however, determining whether this has any relevance for human health has not been possible. Our results demonstrate that functional iKIRs have a biologically significant impact on class II-associated protection, most likely via an impact on class II-restricted protective T cell responses, which manifest as a clinically significant difference in the risk of developing T1D.

## Materials and methods

### Immunogenetic cohorts

The GRID cohort (N=13,452) contains white European individuals from a UK-based case control study. The HBDI collection (Families=402) is a multiplex family dataset comprising families with affected children. The cohorts, ethical approval and the individuals selected for downstream analysis are described in ***SI, Materials and Methods***.

### Statistical analysis

The impact of genotype on disease status was assessed by multiple logistic regression to adjust for covariates. The effect of iKIR score on HLA associations was assessed by stratification and the associated p-values obtained by permutation test. Additional regression approaches and the family-based association analysis are described in ***SI, Materials and Methods***.

### KIR expression analysis

#### KIR gene expression

A published single-cell RNA-seq Case-Control dataset of children progressing to T1D was used to investigate KIR expression variation between cases and controls in NK and T cell populations in blood (EGA study accession: EGAS00001004070). Details on the RNAseq analysis can be found in ***SI, Materials and Methods***.

#### KIR protein expression

To validate the findings obtained with the RNAseq dataset, 20 individuals were recruited (10 T1D cases and 10 controls) for KIR immunophenotyping by flow cytometry (***SI, Materials and Methods***).

## Funding

L.M.B. and C.K. are funded by the European Union H2020 programme under grant agreement 764698 (QUANTII). J.A.T., W.J. and J.T. are funded by European Research Council (ERC) under the European Union’s Horizon 2020 research and innovation programme (grant no. 695551). B.A. is a Wellcome Trust (WT) Investigator (103865Z/14/Z) and is also funded by the Medical Research Council (MRC) (J007439, G1001052), the European Union Seventh Framework Programme (FP7/2007–2013) under grant agreement 317040 (QuanTI), the European Union H2020 programme under grant agreement 764698 (QUANTII) and Leukemia and Lymphoma Research (15012). N.P. is funded by the MRC (1125070). Data from participants in the GRID cohort was kindly provided by J. Inshaw, H. Hong and J. A Todd funded by a Strategic Award to the Diabetes and Inflammation Laboratory from the JDRF (4-SRA-2017-473-A-A) and the Wellcome Trust (107212/A/15/Z).

## Competing interests

The authors declare that they have no competing interests.

## Data and code availability

Scripts used to perform the immunogenetic analysis are available upon request from corresponding author.

## Notes

### Competing Interest Statement

The authors have declared no competing interest.

