## Supplementary Information for "Inhibitory KIRs decrease HLA class II-mediated protection in Type 1 Diabetes"

Supplementary materials and methods.

Supplementary results.

Figs. S1 to S12.

Tables S1 to S16.

Supplementary text.

Supplementary references.

### Supplementary Materials and Methods

#### Subjects

**The GRID case-control cohort.** Our primary cohort is a white European UK-based cohort of 13,452 individuals, with 6,783 cases from the Genetic Resource Investigating Diabetes (GRID) aged between 6 months and 16 years and 6,669 controls from the British 1958 Birth Cohort. Information on disease status was available for GRID individuals. Subjects were genotyped on the Illumina Infinium high density Immunochip array (Illumina Inc., USA). HLA and KIR type were imputed (see below). Individuals with missing information (either sex or HLA/KIR genotype) were discarded, leaving a total of 11,961 individuals (6,219 cases and 5,742 controls).

**The HBDI multiplex family dataset.** Our validation cohort consists of 402 US families from the Human Biological Data Interchange, a repository founded for the purposes of diabetes research[1]. All families have at least 2 affected children and parents are not affected. All 1721 individuals in this repository have been previously genotyped at HLA and KIR genes as described in[2]. Pedigree information was available for all individuals. After removal due to missingness a total of 1700 individuals (342 families) were used in the analysis.

**Cohort for flow cytometry analysis**. We obtained KIR immunophenotype data from 10 T1D patients and 10 matched controls (matching factors: age and gender). Healthy controls age ranged between 22-65 with 60% females. T1D patients had long-standing disease (N=4), intermediate-standing disease (within 3 years of diagnosis, N=1) or were recently diagnosed (within 1 year of diagnosis, N=5). The T1D cohort age ranged between 18-58 with 70% females.

#### Ethical Approval

The immunogenetics component of this study was approved by the Imperial College Joint Research Compliance Office (Ref: 20IC6312). Written informed consent was obtained at the original study sites from all individuals and subjects from both GRID and HBDI cohorts consented to the use of their data in future research studies into the genetic risks of diabetes. For the flow cytometry component (**Cohort for flow cytometry analysis**) all study procedures were conducted according to the principles of the Declaration of Helsinki and all participants gave written informed consent following protocols approved by the relevant ethics committee (NRES London 13/LO/0022, ICREC 21IC7146). Human samples used in this research project were obtained from the Imperial College Healthcare Tissue and Biobank (ICHTB). ICHTB is supported by the National Institute for Health Research (NIHR) Biomedical Research Centre based at Imperial College Healthcare NHS Trust and Imperial College London. ICHTB is approved by Wales REC3 to release human material for research (22/WA/0214).

#### HLA and KIR imputation

HLA alleles at classical HLA loci (*A, B, C, DRB1, DQA1, DQB1* and *DPB1*) were imputed to two field resolution with HIBAG v1.14.0[3]. To ensure high imputation accuracy, call threshold (CT) was set to 0.5 (imputations with posterior probability below CT were not called and left blank). Call rates for classical HLA loci ranged between 92.75% and 99.67%. Imputed genotypes were compared with high-resolution experimentally typed HLA genotypes available in a subset of GRID samples. Prediction accuracy ranged from 86.68% to 98.06% (**Fig. S1A**). For KIR imputation, haplotypes were estimated using SHAPEIT v.r837[4] (parameters: 500 states, 10 burn-in iterations, 50 main iterations, 10 pruning iterations, genetic map HapMap phase II b37). KIR imputation was then performed using KIR*IMP[5] using 231 SNPs at chromosome 19. We kept imputations at a probability threshold of 0.5 which has been reported to ensure 92% accuracy. KIR copy number accuracy was assessed using copy number data on a subset of individuals directly typed by qPCR at *KIR3DL1/S1* locus[6] (**Fig. S1B**).

##### Haplotyping/HLA genotype phasing

Haplotypes at *DRB1* and *DQB1* loci were estimated from the imputed genotypes with the R package Haplo.stats[7]. Haplo.stats uses expectation-maximization algorithm with progressive insertion of loci to estimate the most probable pair of haplotypes for a given genotype. Haplotypes were phased separately in cases and controls.

##### iKIR score and iKIR count

We define a *functional iKIR gene pair* as the presence of both the iKIR gene and the gene encoding its HLA ligand in the same individual[8]. For each individual in the cohort, we calculate the count of functional iKIR genes. We consider the following iKIRs: KIR2DL1, KIR2DL2/KIR2DL3 and KIR3DL1. We also compute the inhibitory score, this is the iKIR count adjusted to reflect the binding strength of the different iKIRs as reported in[8]; replicated here in the **Supplementary text**. We obtain very similar results with both metrics unless stated otherwise in the text. When classifying individuals into “high” or “low” iKIR score strata, the threshold for calling high/low was chosen on purely pragmatic grounds to give balanced numbers in both arms (to minimise problems of sparse data). Multiple thresholds were tested when possible, and conclusions remained unchanged unless stated otherwise.

We also compute an absence of functional iKIR count to investigate whether HLA ligands alone are responsible for the iKIR score effect of HLA associations. This score reflects the number of genes for HLA ligands in the absence of their matching iKIR genes in an individual.

#### Statistical analysis

##### *Logistic Regression*

Multiple logistic regression was used to investigate the effect of iKIRs on the significant HLA associations that we identified in our case-control cohort. We used two regression approaches to assess the impact of the genotype on disease outcome:

**Regression by stratification**. We stratified our cohort into individuals with high and low iKIR score (that is greater than the threshold and less than or equal to the threshold respectively) and then assessed the impact of HLA genotype on outcome using logistic regression within each iKIR stratum (two models). Significance was assessed by a permutation test (see below).

**Regression by interaction.** To explore the role of significant covariates in our cohort (HLA ligands and sex) on the iKIR effect, we modeled the whole cohort and included iKIR score as an interaction term with the HLA genotype of interest in the model $\boldsymbol{OUTCOME \sim HLA\ genotype \times iKIR\_score\ +\ COVARIATES}$.

Significance was given by the p-value of the interaction coefficient *HLAgenotype:iKIRscore*.

##### *Permutation test*

To assess the significance of the difference in odds ratio (OR) between the models in the two strata (high iKIR and low iKIR) for a single class II genotype, we conducted a permutation test using Monte Carlo methods. We permuted (by resampling without replacement) the iKIR score values across all the individuals, stratified the (permuted) cohort and run the models in the two strata. We repeated this process $10^8$ times. Then we counted the number of permutations where the ln[OR] in the iKIR_high strata was equal or greater than the observed ln[OR] in the iKIR_high strata and the ln[OR] in the iKIR_low strata was equal or less than the observed ln[OR] in the low strata. The P-value was the number of permutations with equal or more extreme values than the observed values divided by the total number of permutations. Similarly, to obtain a P-value for the observed difference between strata across all protective HLA class II genotypes associated with T1D, we permuted the iKIR score, calculated the ln[OR] in the iKIR_high strata minus the ln[OR] in the iKIR_low strata and then calculated the mean of these absolute differences across the protective genotypes weighted by the number of genotype-positive T1D cases. We repeated the permutation $10^8$ times and counted the number of iterations where the weighted mean was more extreme than the observed weighted mean in our cohort; the P-value reported is this count of extreme values divided by the total number of permutations.

##### *Identification of genotypes significantly associated with T1D*

To identify genotypes significantly associated with risk of, or protection from, T1D we first identified all alleles at the *HLA-A, -B, -C, -DRB1, -DQA1* and *-DQB1* loci as well as all two and three allele genotypes at *DRB1*, *DQA1* and *DQB1* loci significantly associated with outcome. We considered alleles in cis and in trans as trans-acting effects are well-documented in T1D. At this stage we used a high P-value threshold for significance ($P<5x10^{-4}$) to create an inclusive list. This yielded a list of 21,726 genotypes. Many of these genotypes are highly correlated and neutral genotypes could appear significant because they mark carriage of a causal “driver” genotype. To eliminate these neutral genotypes we asked, for each genotype on the list, whether its protective/detrimental effect was independent of all other genotypes on the list. Specifically, for each genotype we considered it sequentially paired with all other genotypes (a total of 21,696x21,695/2=235,347,360 pairs of genotypes) and assessed whether their protective or detrimental effects were independent of each other by including the two genotypes simultaneously in a multivariate regression and assessing whether i) the direction of their protective/detrimental effect remained unchanged and ii) whether the effect remained significant ($P<5x10^{-4}$). For colinear genotypes we adopted the following rules: if the genotypes were completely colinear (one allele always present with the other) then we retained the longest genotype (as it is impossible to rule out the possibility that the longer genotype was the driving genotype). If the genotypes were weakly colinear we retained the genotype with the lowest P-value in a multiple regression. This yielded a list of 33 alleles or sets of alleles significantly associated with outcome whose effects were independent of all other single alleles or sets of alleles. We then manually curated the list removing two genotypes (*DRB1*07:01-DQA1*01:02* and *DQA1*01:02-DQB1*03:01*) that were compounds of other genotypes on the list and although were independent of each of these genotypes individually were not independent of both of them simultaneously (i.e. in multiple regression with all three genotypes including the eliminated genotype suffered a reversed direction of effect). A third genotype (*DRB1*03:01-DQA1*05:01-DQB1*02:01-DQB1*03:02*) was retained on the list since although it appears to be a compound of the detrimental haplotype *DRB1*03:01-DQA1*05:01-DQB1*02:01* and the detrimental genotype *DQB1*03:02* it retained direction and significance of effect (albeit considerably weakened) in multiple regression when both *DRB1*03:01-DQA1*05:01-DQB1*02:01* and *DQB1*03:02* were included simultaneously with it (presumably due to synergy between the genetic effects). Finally, (having used an inclusive list to make sure alleles were stringently removed due to lack of an independent effect) we then calculated the effective number of tests conducted (for all alleles at the *HLA-A, -B, -C, -DRB1, -DQA1* and *-DQB1* loci as well as all pairs and trios of *DRB1, DQA1* and *DQB1* alleles, taking into account that many of these tests are highly correlated). We considered four methods to estimate the number of effective tests: that of Nyholt, Li and Ji, Gao et al and Galwey[9-12] implemented in the function meff of the poolr package in R. All four methods gave similar values for $M_{eff}$, we took the highest $M_{eff}$ value (obtained with Li & Ji method) to obtain the most stringent cutoff, which via the Bonferroni correction ($P=0.05/M_{eff}$) resulted in a cut off for significance of $P=1.35x10^{-5}$.

##### *Power analysis*

The independent US family dataset was used for validation. This dataset is ten times smaller than the GRID cohort (700 trios vs 6219 cases in the GRID cohort). Therefore, before attempting to validate our results, we used Monte Carlo methods to study the statistical power to detect a significant iKIR score modification given the sample size of our validation set. We assumed that the observed iKIR effect in the GRID cohort was representative of the effect size in the US cohort and used Monte Carlo methods to conduct a simulation-based power analysis. Briefly, we generate a bootstrap subsample (resampling individuals in our cohort with replacement) of the GRID cohort with size equal *s* (Ncontrols=s/2, Ncases=s/2) and we run the regression analysis on this subsample. This process is repeated 100 times. The estimated power for sample size *s* is the proportion of bootstrap subsamples where we retrieved a significant difference across protective genotypes. In this way we assessed which genotypes or combinations of genotypes had sufficient power to permit study in the family dataset.

*Family-based analysis*

We analyzed each family as trios for each affected sibling. Trios were stratified into “KIR high” or “KIR low” based on the iKIR score of the affected child (KIR high: iKIRscore>threshold, KIR low: ≤threshold, thresholds considered: 1.75 in first instance as it yields most balanced division but 1, 1.5, 2 and 2.5 were also considered). Then for both strata and each driver allele, we counted the number of transmissions of the protective allele from heterozygous parents to affected children and the number of non-transmissions i.e. the transmissions of the protective allele to a non-person. Following Spielman et al and Ott[13,14] we calculated b and c in the matrix:

| | | Not transmitted | | |
|---|---|---|---|---|
| | | $M_1$ | $M_2$ | |
| transmitted | $M_1$ | a | b | a+b |
| | $M_2$ | c | d | c+d |
| | | a+c | b+d | 2n |

Where M is the locus of interest, $M_1$ is the protective “driver” allele and $M_2$ all other alleles at that locus.

Our statistic was the difference of the log of the ratio of transmitted to non-transmitted genes between individuals with a high iKIR score and individuals with a low iKIR score i.e

$$\ln[b/c\,|_{KIR\ high}] - \ln[b/c\,|_{KIR\ low}]$$

Whilst Spielman and Ott calculated the distribution of the transmission ratio under their null hypotheses of interest analytically, this approach was not available to us as we were interested in several driver alleles whose transmission could not be guaranteed to be independent of each other. We therefore calculated the null distribution using a Monte Carlo approach. Specifically, the distribution of the statistic under the null hypothesis that iKIR score does not modulate HLA class II associations was assessed by permuting the iKIR score values among affected offspring and then calculating the statistic. This was repeated $10^5$ times. The empirical P-value was the proportion of permutations with an estimate equal or more extreme than the observed estimate.

#### Fraction of cases prevented

Following [15] we define the fraction of cases prevented by a genotype $G^+$ as $F_P$

$$F_P = \frac{P(D|G^-) - P(D)}{P(D|G^-)}$$

which, given the following contingency table

| | $G^+$ | $G^-$ |
|---|---|---|
| D (Disease) | a | b |
| H (Healthy) | c | d |

can be rewritten as

$$F_p = (1-R)\left(1 - \frac{d(a+b)}{b(c+d)}\right)$$

where R is the prevalence of disease.

#### Other genes associated with T1D

We generated a comprehensive list of T1D-associated loci reported to date in the NHGRI-EBI GWAS catalog. The NHGRI-EBI GWAS catalog is a manually curated collection of published GWASs that is regularly updated. Only SNP associations with $P<1x10^{-5}$ from GWASs with more than 100,000 tagged SNPs (preQC) are included in the database.

T1D GWAS-associated SNPs reported in the GWAS catalog were downloaded on the 22nd March 2021 from https://www.ebi.ac.uk/gwas/efotraits/EFO_0001359. The search term *type 1 diabetes* returned a total of 351 associations from 56 different studies. We only kept SNP associations under the EFO trait *type I diabetes mellitus*. The resulting list contained 196 SNP associations from 16 different studies mapping 67 different genes. We only report protein coding genes with significant SNP associations (within the gene or nearby) that have been reported in at least 2 out of the 16 T1D GWAS publications (either the same SNP or different SNPs mapping to the same gene). The mapped genes and their effect sizes are reported together with date when they were first reported. The date of discovery was manually curated to account for the fact that some loci were first reported in candidate gene studies (before GWASs, e.g. HLA region or INS gene).

#### Single-cell RNAseq analysis from peripheral blood mononuclear cell samples

Raw FASTQ files from UMI-based single-cell RNAseq experiment of peripheral blood mononuclear cells were downloaded from the European Genome-Phenome Archive (EGA) with accession number EGAS00001004070. The samples were collected from four children who developed β-cell autoimmunity (i.e. were seropositive for islet auto antibodies) and matched control subjects[16]. Reads were aligned using STARsolo aligner with the option `-soloMultiMappers Uniform`[17]. After read mapping and count quantification, the 8 count matrices were read into R using the Seurat package for quality control[18]. Briefly, empty barcodes, apoptotic or stressed cells and doublets were discarded leaving a total of 18,519 cells. An automated annotation method, SingleR[19], was then used to classify single cells. Given a reference dataset of single-cell or bulk samples with known labels, SingleR classifies new cells from a test dataset based on spearman correlation between the test and the reference. The Monaco and the human primary cell atlas (HPCA) reference datasets from the Celldex package were used[19].

To make feature counts comparable across cells, we used the `NormalizeData()` function from Seurat: feature counts are divided by the cell library size (total umi counts of the cell) and multiplied by 10,000. The resulting feature quantities are natural-log transformed using log1p. So KIR expression values are log normalized KIR counts per 10,000 reads.

#### KIR protein expression analysis

20 individuals including 10 T1D patients and 10 matched controls were recruited for KIR immunophenotyping flow cytometry analysis. The healthy cohort age ranges between 22-65 with 60% females and the T1D cohort age ranges between 18-58 with 70% females. The multi-colour antibody panel (**Table S16**) was designed to assess expression of KIR2DL1, KIR2DL2/L3 and KIR3DL1 in the different CD4+ and CD8+ naive and memory populations as well as in NK cell subsets.

**T cell subsets**

CD28 and CD45RA staining was used to gate CD4+ and CD8+ T cells into the following populations (see **Fig. S9** and **Fig. S10** for representative gating strategy):

- Tnaive: CD28+CD45RA+
- Tcm (central memory): CD28+CD45RA-
- Tem (effector memory): CD28-CD45RA-
- Temra (effector memory expressing CD45RA): CD28-CD45RA+

**NK cell subsets**

CD56 and CD16 staining was used to identify NK cells (see **Fig. S11** for representative gating strategy). First CD56 staining was used to identify NK cells in the CD3-Dump- gate. Next, both CD56 and CD16 markers were used to identify CD56dimCD16+ and CD56brightCD16- NK cell populations.

All samples were stained and analysed on a BD FACSAria III (BD Biosciences) in the same experiment. Automated flow cytometry data analysis was performed in R. `singletGate` method from OpenCyto package was applied to gate out singlets[20]. All remaining gates were based on the mindensity method. Gates were determined using collapsed data across all individuals. Independent manual gating by someone with expertise in flow cytometry gave very similar results.

#### Mathematical model of the transition from health to T1D

We model autoimmune response against beta cells in the pancreatic islet with a set of four coupled differential equations describing the interactions between activated islet specific T cells, both conventional CD8+ (C) and regulatory CD4+ (R) T cells, insulin producing β-cells (B) and islet antigen levels (A):

$$\frac{dB}{dt} = -\delta_B BC \qquad (1)$$

$$\frac{dA}{dt} = \alpha_A BC - \delta_a A \qquad (2)$$

$$\frac{dR}{dt} = \alpha_R A - \delta_T R \qquad (3)$$

$$\frac{dC}{dt} = \alpha_C A - \delta_T C - \delta_i RC \qquad (4)$$

We assume that the presence of islet antigens (A) activates conventional T cells (C) and regulatory T cells (R). We assume that regulatory cells are CD4+, conventional cells could either be CD4+ or CD8+ T cells with effector function. The activation of C and R has two opposite effects; while C mediates β-cell killing, R reduces activated C levels (which in turn reduces β-cell killing). The coexistence of these two antagonistic pathways triggered by the same input constitutes a type of incoherent feedforward loop (IFFL). This motif has been used to describe immune activation and tumor control [21]. The IFFL is coupled to a positive feedback loop: T cell mediated killing of β-cells results in more islet antigen release that in turn activates T cells (C and R).

We explored additional structural forms of the model. We included density-dependent production of T cells and modelled the negative feedback loop via R inhibition on C proliferation:

$$\frac{dB}{dt} = -\delta_{beta} BC \qquad (5)$$

$$\frac{dA}{dt} = \alpha_A BC - \delta_a A \qquad (6)$$

$$\frac{dR}{dt} = \alpha_R AR\left(1 - \frac{R}{K_R}\right) - \delta_T R \qquad (7)$$

$$\frac{dC}{dt} = \frac{\alpha_C AC}{k+R}\left(1 - \frac{C}{K_C}\right) - \delta_T C \qquad (8)$$

We refer to the density-independent and the density-dependent models as model 1 and model 2 respectively. Parameters used for the simulations using each model are listed in **Table S15**.

#### Simulations

We apply the cellular model described above to a virtual cohort of 10,000 individuals, each one defined by a different set of parameters. T cell parameters define the genetic risk of an individual to develop T1D. HLA genes only explain 50% of T1D risk[22] so not all individuals with a susceptible genotype will develop T1D and other genes and factors such as history of infections, microbiome and lifestyle are thought to contribute to the transition to T1D. To overcome these unknowns, we simulate a cohort of young seropositive individuals, i.e., individuals that are positive for islet antigens. This way we have a homogeneous population of individuals of similar age with an active adaptive response (triggered by an enteroviral infection for example). Even though seropositivity for islet antigens is associated with T1D, not all seropositive individuals progress to overt T1D[23]. Thus, depending on each individual parameters, the immune response will be resolved without significant β-cell loss or alternatively will lead to massive β-cell killing, resulting in T1D onset. We assume similar dynamics (on average) in all the pancreatic islets and that loss of 80% of the beta cells in the islet leads to T1D.

#### Software

All calculations were performed using custom scripts in R (v4.0.3). Packages used include stats (v3.6.2), parallel (v4.0.3), data.table (v1.13.2), Haplo.stats (v1.8.6), poolr (v0.8-2) and debug (v1.3.83).

### Supplementary Results

#### iKIR modification of *DRB1*15:01-DQB1*06:02*

In the main text we focus on whether iKIR modify the *DQA1*01:02-DQB1*06:02* association as there is some evidence in the literature[24,25] and in our cohort that the protection associated with the *DRB1*15:01-DQA1*01:02-DQB1*06:02* genotype maps to *DQA1*01:02-DQB1*06:02* rather than *DRB1*15:01-DQB1*06:02*. However, fine mapping is difficult, so in case the driver of protection is *DRB1*15:01-DQB1*06:02* we repeated our iKIR analysis focusing on *DRB1*15:01-DQB1*06:02* instead. As expected, (since 99.2% of *DQA1*01:02-DQB1*06:02* positive carriers are also *DRB1*15:01-DQB1*06:02*), the results are virtually identical (**Fig. S6-S7**, **Table S8-S11**).

#### Modification of *DQA1*01:02-DQB1*06:02* is attributable to functional iKIR rather just the HLA class I genes

An individual's iKIR score will depend on their HLA class I genotype since the iKIR ligands are HLA class I molecules. Some HLA class I alleles are associated with protection from or susceptibility to T1D[26-28]. Therefore, our observation that *DQ6* protection is enhanced in individuals with a low iKIR score could be due to an interaction between HLA class I alleles and *DQ6* and have nothing to do with iKIR genes. To investigate whether our observations were attributable to the class I ligands effects rather than the iKIR genes, we repeated all the analysis with iKIR ligands (Bw4, C1, C2) included as covariates. The results were remarkably similar and iKIR score still significantly modified the *DQ6* protective effect even when iKIR ligands were included as covariates (**Table S12**).

To further investigate whether the observed modification of *DQ6* was attributable to the effects of class I ligands rather than the iKIR genes, we constructed a score to reflect presence of the HLA ligand but absence of the matching iKIR. Although this score in itself was associated with significant protection (ln OR = -0.4 P =0.0009) in line with the published literature that HLA class I molecules are associated with risk of T1D, it had no impact on the protective effect of *DQ6* (P=0.99) when included as an interaction term ($OUTCOME \sim DQ6 \times ligand\_no\_iKIR + GENDER$) suggesting that the effect we had observed for iKIR score (functional iKIRs) requires presence of the gene encoding the iKIR not just the HLA ligand. However, we were concerned that there was little variation in the count of ligands in absence of iKIR since most individuals have at least one functional iKIR in our cohort. This could have reduced the power of the test resulting in the non-significant result, so we also developed the "ligand count" measure which quantified the number of HLA ligands. This is a more problematic measure as it is significantly correlated with the iKIR score (corr=0.87, $P<2x10^{-16}$) making it difficult to disentangle the effects. However, we found that iKIR score has a stronger effect than ligand count on *DQ6* ($P=5.5x10^{-6}$ for iKIR score, $P=7.6x10^{-4}$ for ligand count); in backwards stepwise regression (starting from a full model with both interaction terms) ligand count is removed from the model and the model with ligand count has a considerably lower AIC (difference=75). Upon standardising the variables for comparability, we also found iKIR score had a higher coefficient than the ligand count (0.62 for iKIR score 0.45 for ligand count). Additionally, in a model including both interaction terms, iKIR score remained significant ($P= 9x10^{-4}$) whilst HLA ligand count became non-significant (P=0.29).

Finally, we assessed whether the iKIR score effect on *DQ6* was attributable to specific HLA class I drivers in our cohort associated with this haplotype. A detrimental Bw4 or a protective Bw6 allele in linkage disequilibrium with *DQ6* could potentially explain the iKIR score effect on *DQ6* since for example people with the detrimental Bw4 allele would tend to have a higher iKIR score than those without (being in possession of an iKIR ligand) and this detrimental effect could act to reduce the protective effect of *DQ6* in the iKIR low strata. Likewise, a protective Bw4 or a detrimental Bw6 allele negatively associated with *DQ6* could also be responsible for the *DQ6*:*iKIR_score* effect. If a particular HLA-B allele was responsible for the iKIR effect, then on removing individuals with this allele the iKIR score effect will disappear. We tested this hypothesis by discarding all individuals with a given allele and then performing regression analysis in the resulting subcohort. We repeated this process for all HLA class I alleles, one allele at a time. For all the HLAclassI_allele$^{negative}$ subcohorts, the *DQ6*:*iKIR_score* interaction term remained significant (**Table S13**).

Overall, this set of results indicate that the observed functional iKIR modification of *DQ6* is not driven by HLA class I genes alone and suggests that the effect of functional iKIR on this protective genotype is due to the iKIR-ligand interaction.

##### All functional iKIRs contribute to iKIR score modulation of *DQ6*

In a logistic regression model, the *DQ6*:Bw4 interaction term is significant (P=3.67x10$^{-4}$), although iKIR score as an interaction term is more significant (P=2.1x10$^{-5}$). Since 95% of Bw4 carriers carry the corresponding iKIR gene, *KIR3DL1*, we hypothesized that the *DQ6*:Bw4 interaction was reflecting the effect of *KIR3DL1-Bw4* gene pair on *DQ6* and that perhaps the iKIR score effect was only driven by functional *KIR3DL1*. Therefore, we investigated the contribution of the rest of iKIR-ligand pairs to the iKIR score by excluding non-functional *KIR3DL1* individuals from the cohort (so that the cohort was homogeneous for functional *KIR3DL1*) and then repeating the stratification analysis on the resulting subcohort. We obtained similar results despite the much smaller cohort size (**Fig. S8**). Similarly, *DQ6:iKIR_score* interaction term remains significant in this subcohort (ln[OR]=1.2, $P=2.53x10^{-4}$) suggesting that KIR2DL1/KIR2DL2/KIR2DL3 also contribute to the modulatory effect along with KIR3DL1.

#### Are activating KIRs modulating HLA associations?

Some studies have reported an association between activating KIRs (aKIRs) and increased risk of T1D[29,30]. We therefore investigated whether the observed iKIR effect could be explained by activating KIR genes. We computed a measure similar to the iKIR count but for the functional activating KIRs, the aKIR Count, including all activating KIR with known ligands namely KIR2DS1, KIR2DS2, KIR2DS4 and KIR3DS1[31]; lack of information about the strength of signaling of these ligand-aKIR interactions precluded development of a score for activating KIR. Although the iKIR score is significantly correlated with aKIR count (corr=0.65, $P<2.2x10^{-16}$), in a model including both metrics interacting with *DQ6*, iKIR score remained significant whereas aKIR count did not.

#### iKIR modulation is independent of detrimental class II haplotypes

The most significant detrimental genotypes in T1D are *DR3* (*DRB1*03:01-DQB1*02:01*) and *DR4* (*DRB1*04:01/02/04/05-DQB1*03:02*)[32]. We investigated whether the iKIR score effect on *DQ6* varied when these genotypes were included as covariates. We consistently found that, whilst the *DR3* and *DR4* genotypes were highly detrimental they had little or no impact on the interaction between iKIR score and *DQ6*; that is both the coefficient of the interaction (for standardised score) and the P-value were very similar with or without inclusion of the detrimental haplotypes (**Table S14**). This indicates that the impact of iKIR on *DQ6* is independent of the strong detrimental genes of the *DR3* and *DR4* genotypes.

### Supplementary Figures

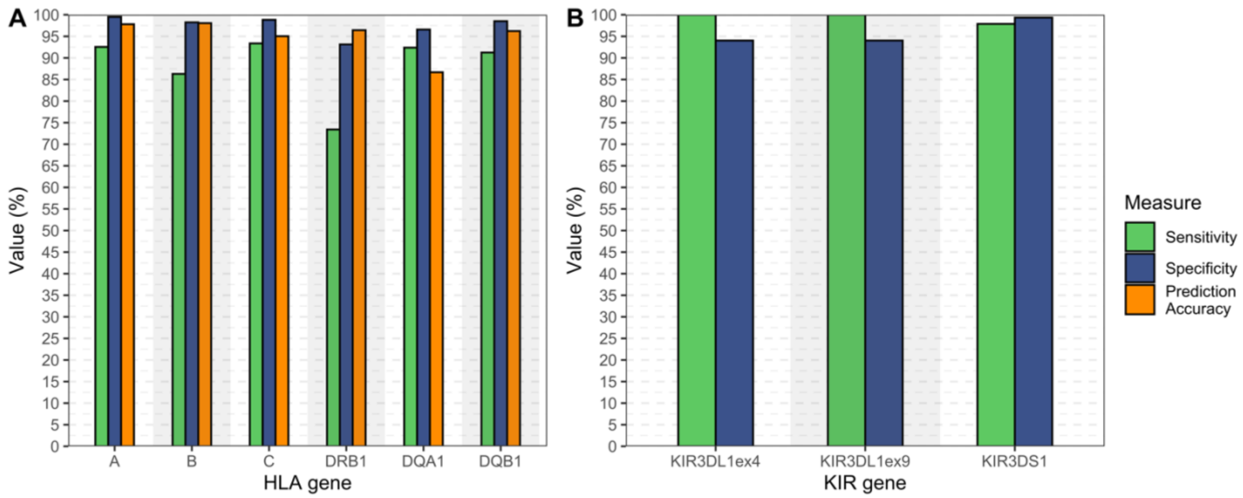

**Fig. S1. HLA and KIR imputation accuracy. A** Imputation accuracy measures at each HLA loci. Prediction accuracy is computed as the number of correctly imputed alleles divided by the number of experimentally typed chromosomes. **B** Sensitivity and specificity values at *KIR3DL1*/*S1* locus. *KIR3DL1* CN was measured using two assay methods, one targeting *KIR3DL1* exon 4 (KIR3DL1ex4) and the other assay targeting exon 9 (KIR3DL1ex9).

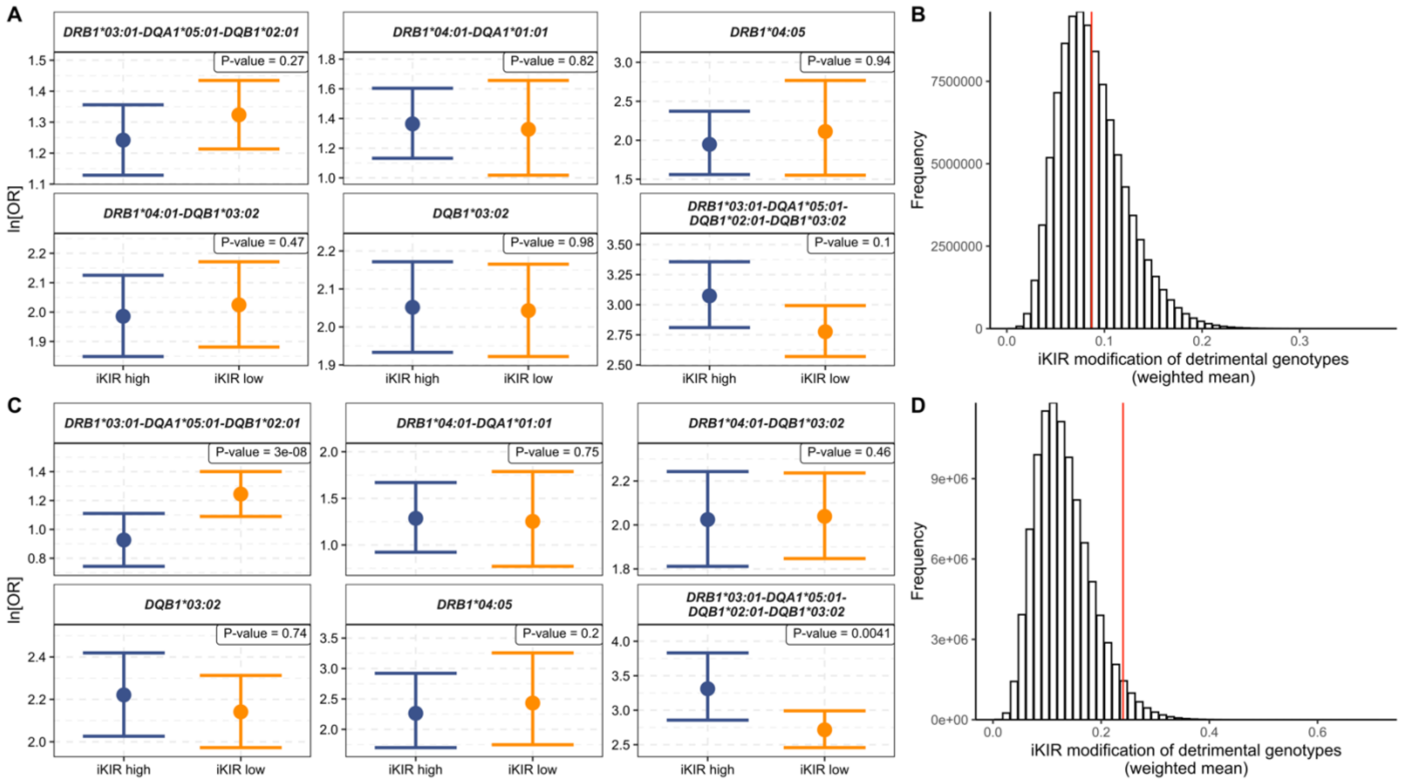

**Fig. S2. Functional iKIR do not consistently modify detrimental genotypes. A** In the whole cohort, for most detrimental genotypes, iKIR have no impact on the detrimental effect (the ln[OR] of the detrimental genotype is very similar in the strata with a high iKIR_score (blue) and the strata with a low iKIR_score (orange) and the 95% confidence intervals are overlapping). Two genotypes (*DRB1*03:01-DQA1*05:01-DQB1*02:01* and *DRB1*03:01-DQA1*05:01-DQB1*02:01-DQB1*03:02*) show some evidence of iKIR modification but the modifications are in opposite directions. **B** The observed value of our test statistic (weighted mean of the difference in ln[OR] between the KIR high and the KIR low strata at threshold=1.75), indicated by the red line, is entirely consistent with the distribution (grey histogram) of the same test statistic under the null hypothesis that the iKIR score has no impact on the detrimental genotypes (generated by permuting the iKIR score of individuals in the cohort). This indicates that the probability of obtaining our observation by chance is high (P=0.46) and that there is no evidence to reject the null hypothesis of no iKIR modification. **C** Similarly in the cohort with HLA class I drivers removed. Most genotypes are not modified and where there is modification then results are in opposite directions. **D**. In the cohort without HLA class I drivers the observed value of our test statistic (indicated by the red line), whilst still overlapping, is more of an outlier from the distribution (grey histogram) of the same test statistic under the null hypothesis that the iKIR score has no impact on the detrimental genotypes. This decrease in the test statistic is driven entirely by one genotype which is strongly iKIR modified (*DRB1*03:01-DQA1*05:01-DQB1*02:01*). Overall, there is some evidence for iKIR modification in this cohort (P=0.04) but it is far from convincing. Examining the apparent iKIR modification of *DRB1*03:01-DQA1*05:01-DQB1*02:01* we found that it was explained by the negative correlation of *DRB1*03:01-DQA1*05:01-DQB1*02:01* with the protective genotype *DQA1*01:02-DQB1*06:02*, which is modified by functional iKIR score since, upon exclusion of *DQB1*0602* carriers, *DRB1*03:01-DQA1*05:01-DQB1*02:01* was no longer iKIR modified (the converse was not the case i.e. *DQA1*01:02-DQB1*06:02* was still significantly modified by functional iKIR when *DRB1*03:01-DQA1*05:01-DQB1*02:01* carriers were excluded from the cohort). In the corresponding permutation test the p-value becomes even less significant (P=0.19). We conclude that there is no evidence that detrimental class II drivers are modified by functional iKIR.

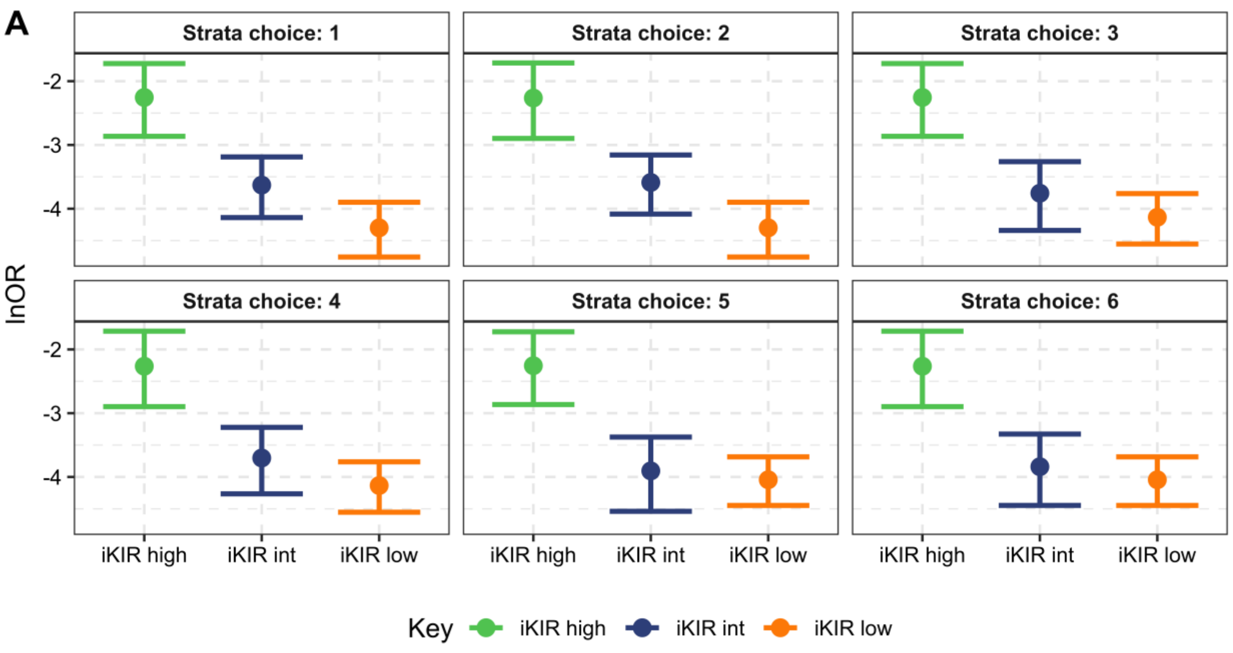

**B**

| Strata choice | Low iKIR category | Intermediate iKIR category | High iKIR category |
|---|---|---|---|
| 1 | [0,1.75] N= 22 | (1.75,2.75] N= 18 | (2.75,4] N= 14 |
| 2 | [0,1.75] N= 22 | (1.75,3] N= 19 | (3,4] N= 13 |
| 3 | [0,2] N= 26 | (2,2.75] N= 14 | (2.75,4] N= 14 |
| 4 | [0,2] N= 26 | (2,3] N= 15 | (3,4] N= 13 |
| 5 | [0,2.5] N= 28 | (2.5,2.75] N= 12 | (2.75,4] N= 14 |
| 6 | [0,2.5] N= 28 | (2.5,3] N= 13 | (3,4] N= 13 |

**Fig. S3. *DQ6* protection as a function of iKIR score**. **A** DQ6 protection increases (i.e. ln[OR] becomes more negative) as the iKIR score decreases. **B** This was true for all strata choices (definitions of high, intermediate, low) considered as shown in the table. Subjects were categorized as having low, intermediate (int) and high iKIR score using all category cutoffs that ensured more than 12 individuals in each group. Coefficients, p-values and group sizes are reported in **Table S2**.

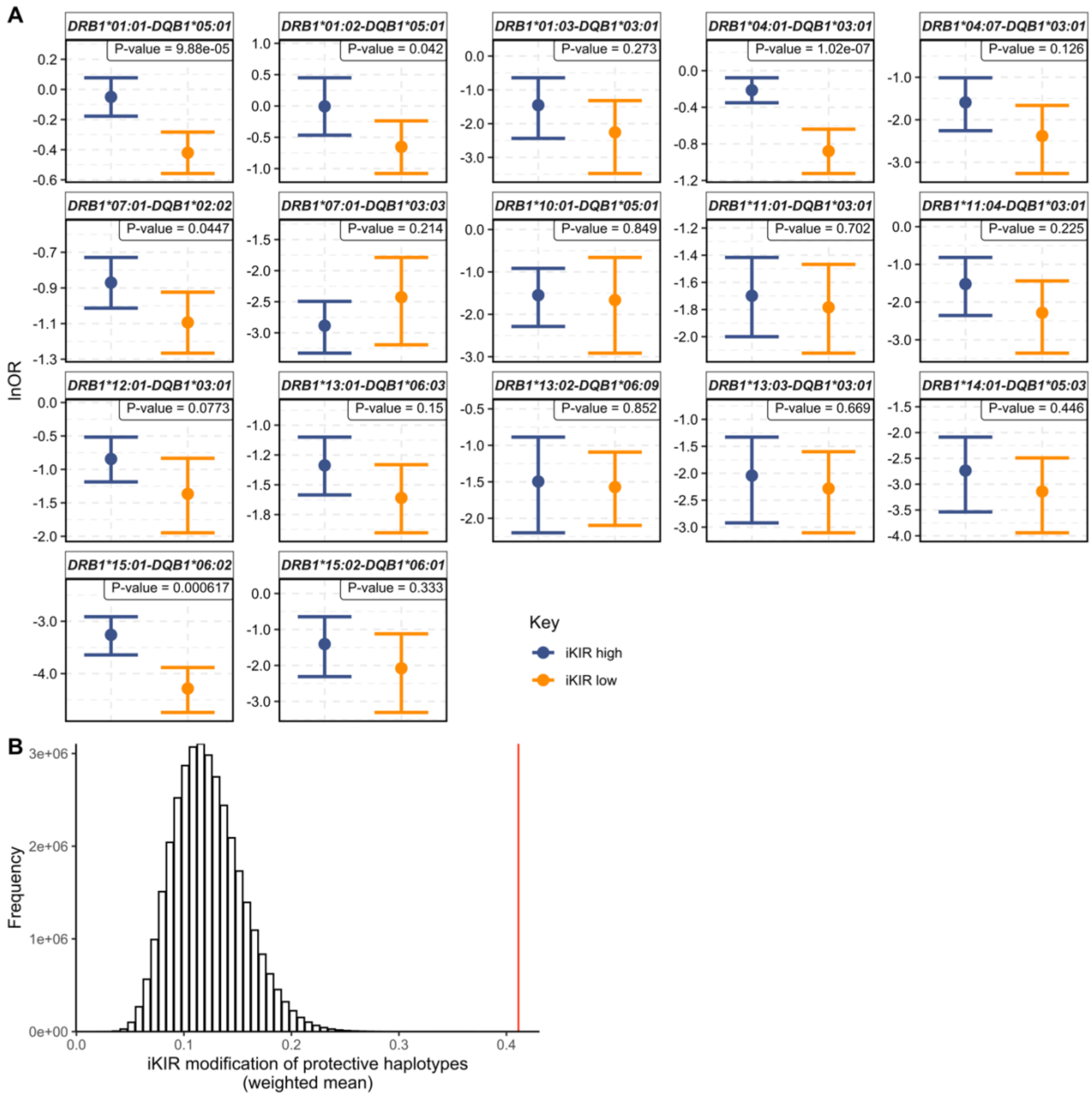

**Fig. S4. Impact of iKIR score on 17 protective *DRB1-DQB1* haplotypes. A** The protective effect of class II *DRB1-DQB1* haplotypes is enhanced in the group of individuals with an iKIR score equal to 1.75 or lower (iKIR score threshold = 1.75) with the exception of *DRB1*07:01-DQB1*03:03*. The number in the top right box corresponds to the odds of seeing this difference by chance ($3x10^7$ permutations). The dot is the ln[OR] and the bars the 95% confidence intervals obtained from the regression, red iKIR high strata, blue: iKIR low strata. **B** The observed value of our test statistic (weighted mean), indicated by the red arrow, lies far above the distribution (grey histogram) of the same test statistic under the null hypothesis that the iKIR score has no impact on the protective haplotypes (generated by permuting the iKIR score of individuals in the cohort, $P<3x10^{-7}$).

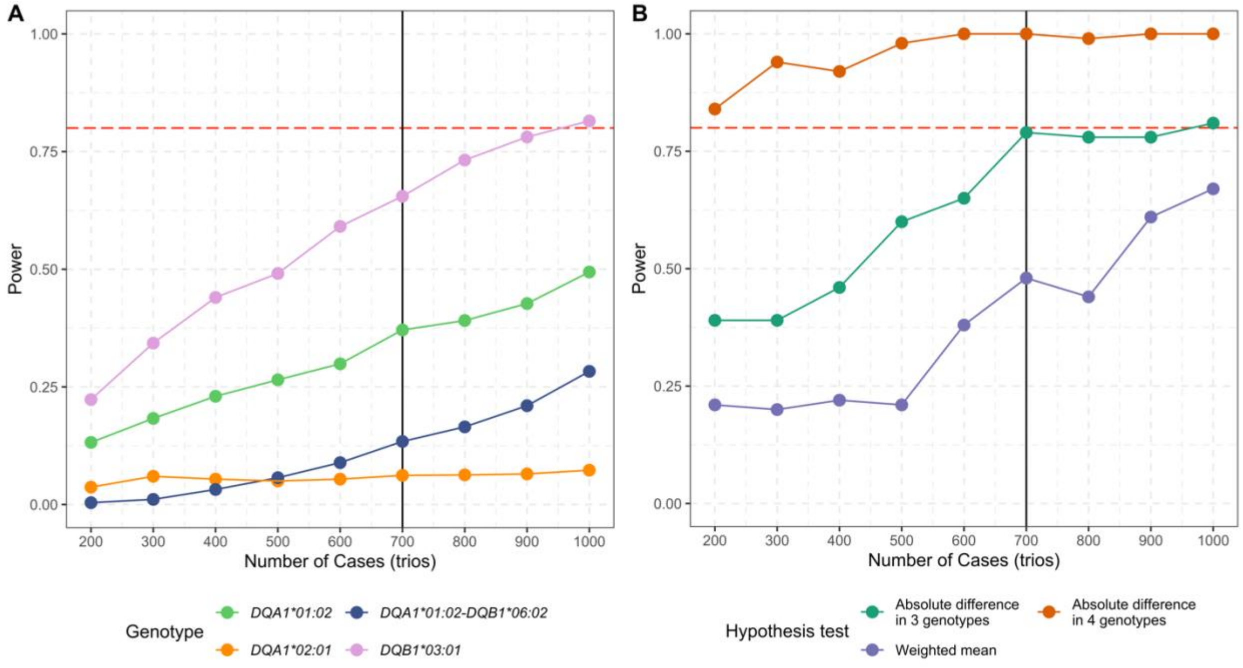

**Fig. S5. Simulation-based power analysis in the GRID cohort. A** For each sample size s (Number of cases=Number of controls=s/2), we generated 1000 random subcohorts by resampling with replacement individuals from the GRID cohort. The sample size of the family dataset is 700 trios (indicated by a black line) which is equivalent to a case control dataset consisting of 700 cases and 700 controls. For each protective genotype and each subcohort we run a logistic regression model with stratified iKIR score (threshold=1.75) as an interaction term with the protective genotype. Power is estimated as the number of subcohorts where the coefficient of interaction was significant (P<0.05). The dashed red line indicates the power standard of 0.8. Given the sample size of the family dataset, the power to detect a significant iKIR score interaction on individual protective genotypes falls considerably below the recommendation of 0.8. **B**. For each sample size, we estimated the power to detect significant difference between iKIR high and iKIR low strata (assessed by permutation test) across several protective genotypes simultaneously. Power is calculated as the proportion of cohorts with a significant permutation test. Given the sample size of the family dataset, the power of detecting an iKIR difference across

4 frequent protective genotypes i.e., *DQA1*01:02*, *DQB1*03:01*, *DQA1*02:01* and *DQA1*01:02-DQB1*06:02* is higher than the recommended standard of 0.8. Sample size family dataset (vertical black line), recommended power (horizontal dashed red line).

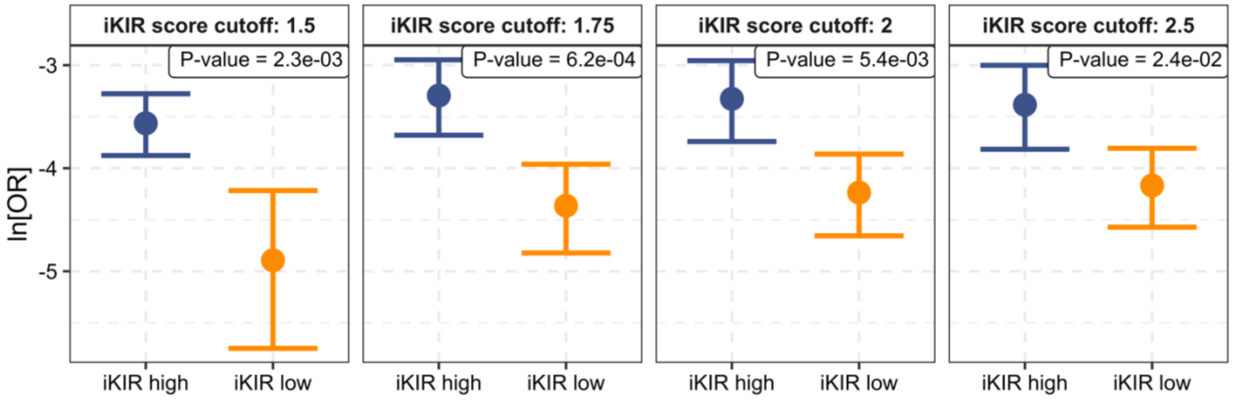

**Fig. S6.** ***DRB1*15:01-DQB1*06:02*** **associated protection is enhanced amongst individuals with low number of functional iKIR.** Stratification analysis was repeated for different iKIR thresholds but this time including ligands as covariates in the model ($\boldsymbol{OUTCOME} \sim \boldsymbol{DRB1*15:01} - \boldsymbol{DQB1*06:02} + \boldsymbol{GENDER} + \boldsymbol{Bw4} + \boldsymbol{C1} + \boldsymbol{C2}$). The results are remarkably similar to our previous analysis on *DQ6* (see main text **Fig. 1**). Estimates, p-values and cohort sizes are reported in **Table S6**.

A

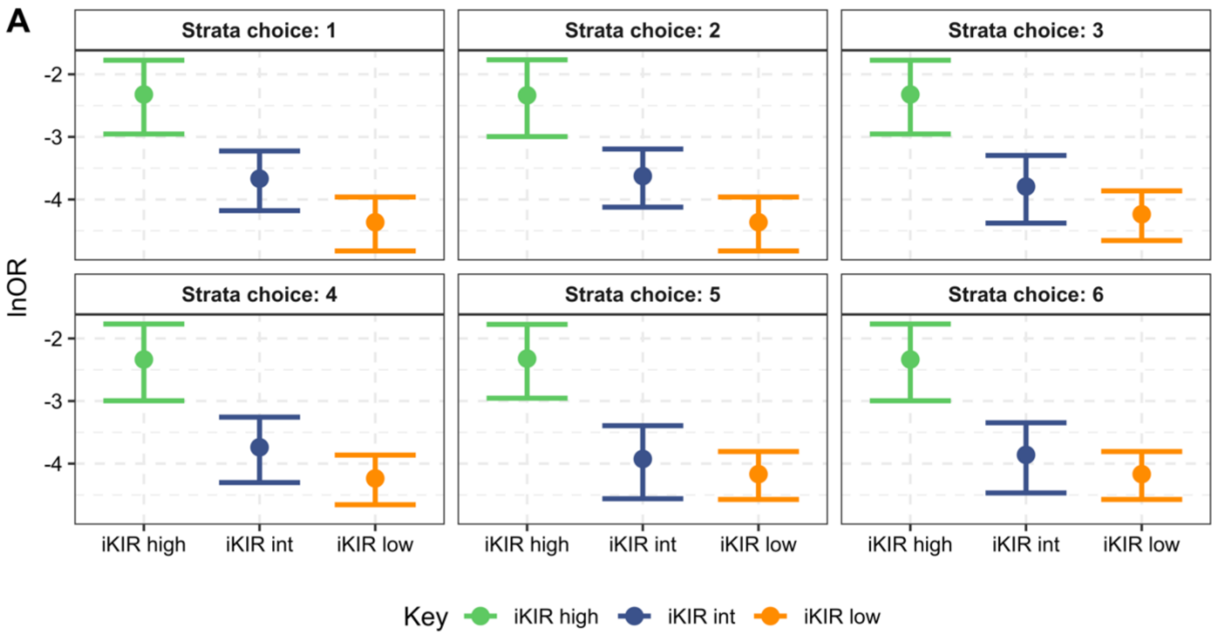

B

| Strata choice | Low iKIR category | Intermediate iKIR category | High iKIR category |
|---|---|---|---|
| 1 | [0,1.75] N= 22 | (1.75,2.75] N= 18 | (2.75,4] N= 13 |
| 2 | [0,1.75] N= 22 | (1.75,3] N= 19 | (3,4] N= 12 |
| 3 | [0,2] N= 26 | (2,2.75] N= 14 | (2.75,4] N= 13 |
| 4 | [0,2] N= 26 | (2,3] N= 15 | (3,4] N= 12 |
| 5 | [0,2.5] N= 28 | (2.5,2.75] N= 12 | (2.75,4] N= 13 |
| 6 | [0,2.5] N= 28 | (2.5,3] N= 13 | (3,4] N= 12 |

**Fig. S7. *DRB1*15:01-DQB1*06:02* associated protection as a function of iKIR score**. **A** The ln[OR] of *DRB1*15:01-DQB1*06:02* decreases (i.e. becomes more protective) as the iKIR score decreases. **B** This was true for all strata choices (i.e. definitions of high, intermediate, low) considered as shown in the table (see also **Table S9**). Subjects were categorized as having low, intermediate (int) and high iKIR score using different thresholds that ensured enough number of individuals in each group (at least N=12). Here, ligands (Bw4, C1 and C2) were included in the model as covariates. These results for *DRB1*15:01-DQB1*06:02* are very similar to the results for *DQA1*01:02-DQB1*06:02* (see **Fig. S3**).

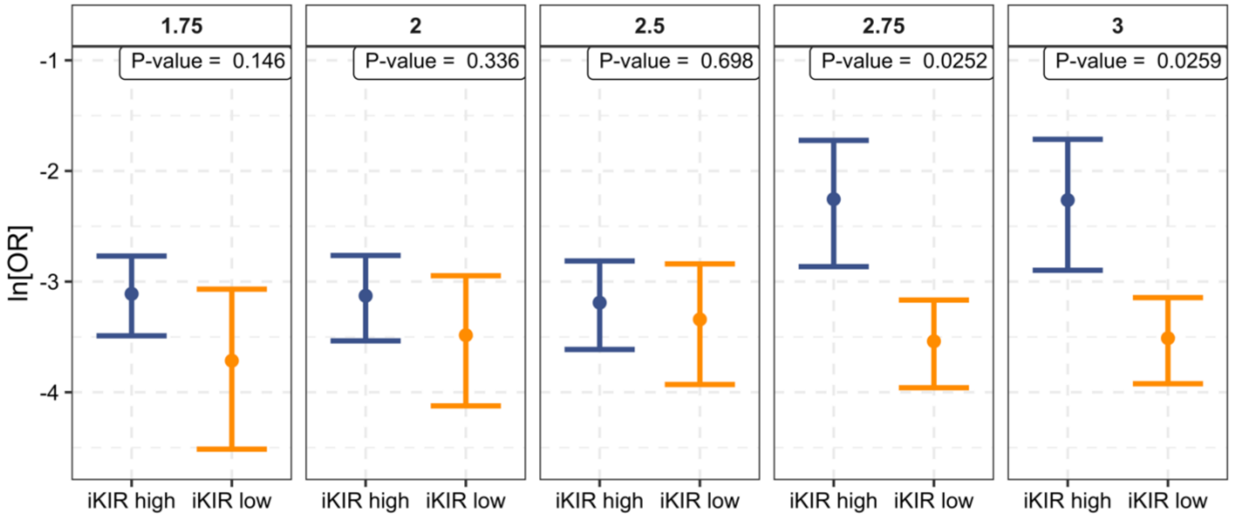

**Fig. S8. iKIRs other than KIR3DL1 contribute to the iKIR score effect on *DQ6*.** In a subcohort in which all individuals carry functional KIR3DL1 we still observe an enhanced protection of *DQ6* in individuals with low iKIR score. The number on the top right box corresponds to the odds of seeing this difference by chance ($10^8$ permutations).

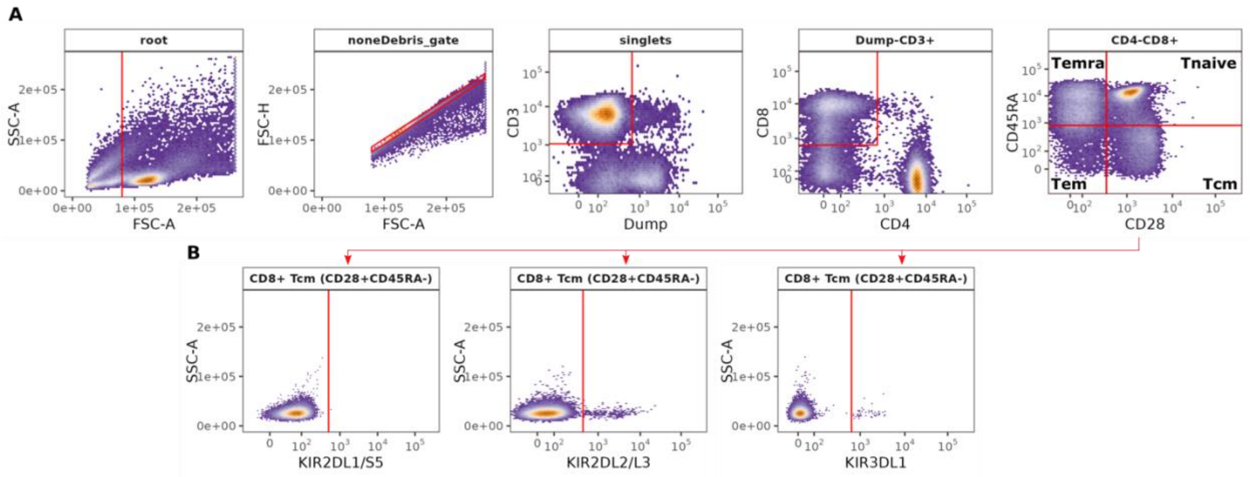

**Fig. S9. Gating strategy for enumeration of iKIR+ CD8+ T cell subsets.** Analysis of PBMCs for a healthy donor (LD1). Each subplot shows events in the parent population, where strip names indicate parent population (root=all events). **A** From left to right, serial gating is used to identify (1) lymphocytes and discard debris (root, all events), (2) single cells in the nonDebris gate, (3) CD3+ and Dump- cells in the singlet gate (excluding unwanted lineages like CD14, CD19 and also necrotic cells), (4) CD8+ T cells in the Dump-CD3+ gate and (5) naive (Tnaive), central memory (Tcm), effector memory (Tem) and effector memory RA+ (Temra) populations within the CD8+ gate using CD45RA and CD28 staining. **B** Each of the 4 subsets defined in the CD8+ gate (naive and memory subsets) was gated to determine events positive for KIR2DL1 (left), KIR2DL2/L3 (middle) and KIR3DL1 (left). In this case, only KIR gates within the Tcm population are shown but the same strategy is followed for Tem, Temra and Tnaive subsets. All boundaries are determined with the 1D mindensity function (except for singlets) using collapsed data across all individuals.

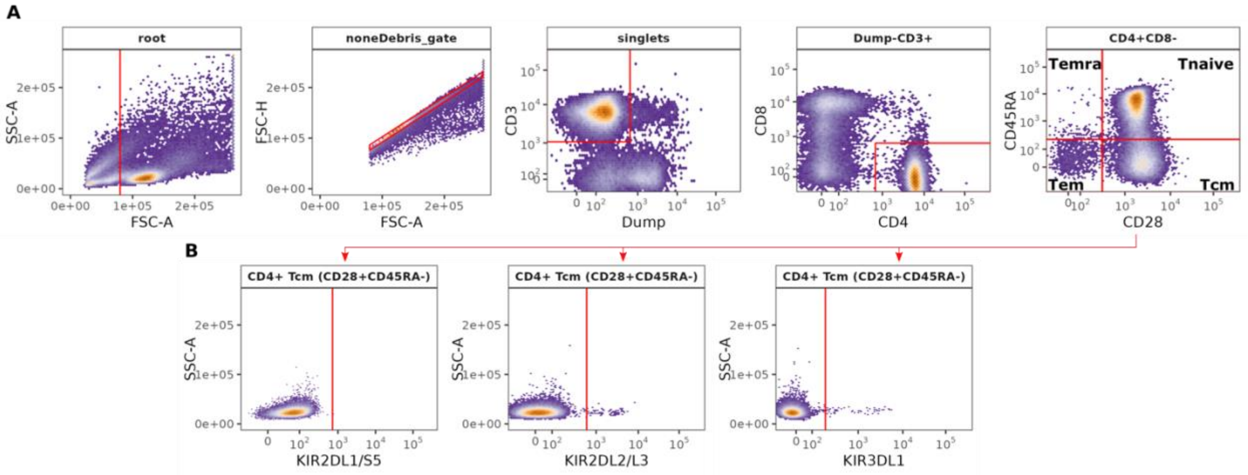

**Fig. S10. Gating strategy for enumeration of iKIR+ CD4+ T cell subsets.** Analysis of PBMCs for a healthy donor (LD1). Each subplot shows events in the parent population, where strip names indicate parent population (root=all events). **A** From left to right, serial gating is used to identify (1) lymphocytes and discard debris (root, all events), (2) single cells in the nonDebris gate, (3) CD3+ and Dump- cells in the singlet gate (excluding unwanted lineages like CD14, CD19 and also necrotic cells), (4) CD4+ T cells in the Dump-CD3+ gate and (5) naive (Tnaive), central memory (Tcm), effector memory (Tem) and effector memory RA+ (Temra) populations within the CD84+ gate using CD45RA and CD28 staining. **B** Each of the 4 subsets defined in the CD4+ gate (naive and memory subsets) was gated to determine events positive for KIR2DL1 (left), KIR2DL2/L3 (middle) and KIR3DL1 (left). In this case, only KIR gates within the Tcm population are shown but the same strategy is followed for Tem, Temra and Tnaive subsets. All boundaries are determined with the 1D mindensity function (except for singlets) using collapsed data across all individuals.

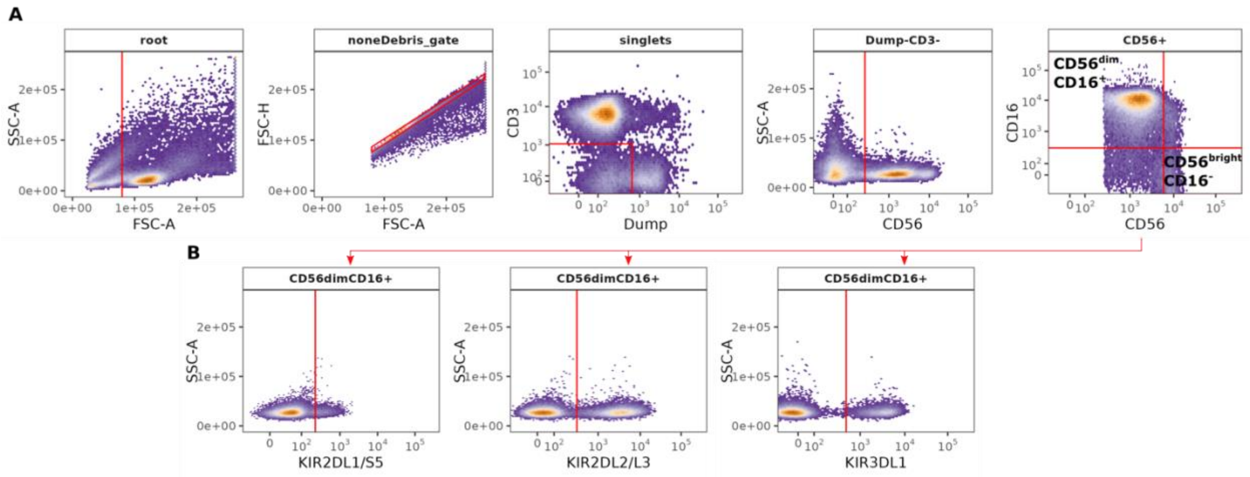

**Fig. S11. Gating strategy to enumerate of iKIR+ NK cell subsets.** Analysis of PBMCs for a healthy donor (LD1). Each subplot shows events in the parent population, where strip names indicate parent population (root=all events). **A** From left to right, serial gating is used to identify (1) lymphocytes, (2) single cells, (3) CD3- cells (CD3-Dump-), (4) NK cells using CD56 staining and (5) CD56dimCD16+ and CD56brightCD16- populations using CD56 and CD16 staining. **B** CD56dimCD16+ and CD56brightCD16- populations were gated to determine events positive for KIR2DL1 (left), KIR2DL2/L3 (middle) and KIR3DL1 (left). In this case, only KIR gates within the CD56dimCD16+ population are shown. All boundaries are determined with the 1D mindensity function (except for singlets) using collapsed data across all individuals.

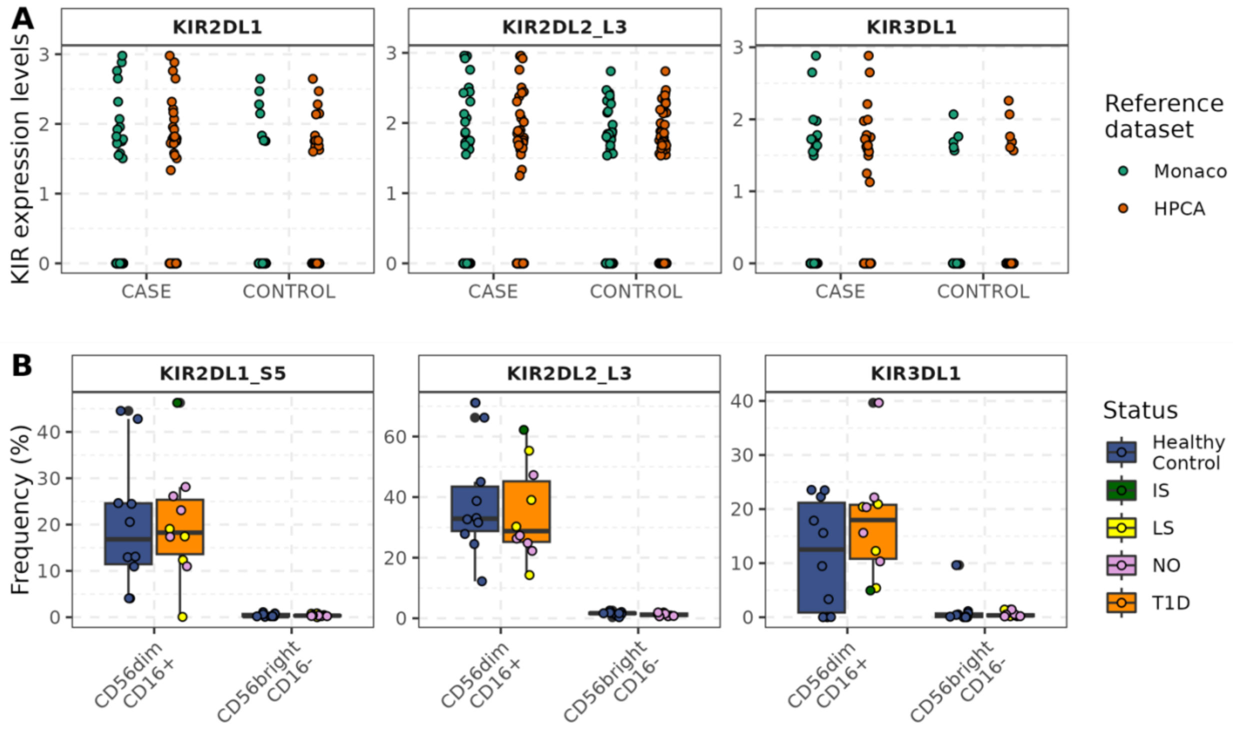

**Fig. S12. KIR+ NK cells are not increased in T1D**. **A** *KIR* gene expression in NK cells split by disease status (CASE=seropositive individuals, CONTROL= matched healthy individuals) and by reference dataset used for cell annotation (green=Monaco reference, orange=Human Primary Cell Atlas reference). Each dot indicates a single cell barcode. Cells labelled as *Natural killer cells* (Monaco reference) or *NK_cells, NK_cell:CD56hiCD62L+*, *NK_cell:IL2* (Human Primary Cell Atlas reference) are shown. **B** Percentage of NK cells expressing different iKIR in T1D patients and healthy controls. The percentage of KIR+ cells in each NK cell population (CD56dimCD16+ and CD56brightCD16-) was quantified by flow cytometry. Dots represent cell frequencies from parent population for each individual. T1D samples are colour coded according to disease duration at time of collection (NO=new onset, IS=intermediate standing disease, LS=long standing disease). Boxes show medians and interquartile ranges within T1D individuals (N=10, orange, irrespective of disease duration) and healthy individuals (N=10, blue).

### Supplementary Tables

| | Group | ln[OR] | 2.50% | 97.50% | P-value | N genotype + | | N genotype- | |
|---|---|---|---|---|---|---|---|---|---|
| | | | | | | Cases | Controls | Cases | Controls |
| | Whole cohort | -3.741 | -4.026 | -3.477 | $1.05\text{x}10^{-157}$ | 54 | 1545 | 6165 | 4197 |
| Threshold=1.5 | iKIR high | -3.49 | -3.79 | -3.20 | $1.82\text{x}10^{-3}$ | 47 | 1165 | 4692 | 3573 |
| | iKIR low | -4.85 | -5.71 | -4.18 | | 7 | 380 | 1473 | 624 |
| Threshold=1.75 | iKIR high | -3.23 | -3.61 | -2.89 | $5.6\text{x}10^{-4}$ | 32 | 721 | 2771 | 2463 |
| | iKIR low | -4.30 | -4.76 | -3.90 | | 22 | 824 | 3394 | 1734 |
| Threshold=2.0 | iKIR high | -3.26 | -3.67 | -2.90 | $4.62\text{x}10^{-3}$ | 28 | 629 | 2400 | 2067 |
| | iKIR low | -4.13 | -4.55 | -3.76 | | 26 | 916 | 3765 | 2130 |
| Threshold=2.5 | iKIR high | -3.33 | -3.75 | -2.95 | $1.87\text{x}10^{-2}$ | 26 | 611 | 2267 | 1911 |
| | iKIR low | -4.04 | -4.45 | -3.68 | | 28 | 934 | 3898 | 2286 |

**Table S1. *DQ6* protection in T1D is enhanced amongst individuals with a low iKIR score.** The cohort was stratified into individuals with high or low iKIR score using different iKIR score thresholds (1.5, 1.75, 2.0 and 2.5). The protective effect of DQ6 was evaluated independently in each stratum using multivariate logistic regression with gender included as a covariate. DQ6 is significantly protective (ln[OR]= -3.74, P=1.05x10-157) in the GRID cohort (Group=Whole cohort, unstratified analysis). Regression coefficients, p-values and cohort sizes are reported for the different strata. P-value for the unstratified analysis calculated using Wald-test, all other p-values calculated using a permutation test.

| Strata choice | Group | | ln[OR] | 2.50% | 97.50% | N genotype+ | | N genotype- | |
|---|---|---|---|---|---|---|---|---|---|
| | | | | | | Cases | Controls | Cases | Controls |
| 1 | High | (2.75,4] | -2.26 | -2.86 | -1.72 | 14 | 126 | 513 | 480 |
| | Int | (1.75,2.75] | -3.63 | -4.14 | -3.19 | 18 | 595 | 2258 | 1983 |
| | Low | [0,1.75] | -4.30 | -4.76 | -3.90 | 22 | 824 | 3394 | 1734 |
| 2 | High | (3,4] | -2.26 | -2.90 | -1.71 | 13 | 120 | 469 | 446 |
| | Int | (1.75,3] | -3.59 | -4.08 | -3.16 | 19 | 601 | 2302 | 2017 |
| | Low | [0,1.75] | -4.30 | -4.76 | -3.90 | 22 | 824 | 3394 | 1734 |
| 3 | High | (2.75,4] | -2.26 | -2.86 | -1.72 | 14 | 126 | 513 | 480 |
| | Int | (2,2.75] | -3.76 | -4.34 | -3.26 | 14 | 503 | 1887 | 1587 |
| | Low | [0,2] | -4.13 | -4.55 | -3.76 | 26 | 916 | 3765 | 2130 |
| 4 | High | (3,4] | -2.26 | -2.90 | -1.71 | 13 | 120 | 469 | 446 |
| | Int | (2,3] | -3.70 | -4.27 | -3.22 | 15 | 509 | 1931 | 1621 |
| | Low | [0,2] | -4.13 | -4.55 | -3.76 | 26 | 916 | 3765 | 2130 |
| 5 | High | (2.75,4] | -2.26 | -2.86 | -1.72 | 14 | 126 | 513 | 480 |
| | Int | (2.5,2.75] | -3.90 | -4.54 | -3.37 | 12 | 485 | 1754 | 1431 |
| | Low | [0,2.5] | -4.04 | -4.45 | -3.68 | 28 | 934 | 3898 | 2286 |
| 6 | High | (3,4] | -2.26 | -2.90 | -1.71 | 13 | 120 | 469 | 446 |
| | Int | (2.5,3] | -3.84 | -4.45 | -3.33 | 13 | 491 | 1798 | 1465 |
| | Low | [0,2.5] | -4.04 | -4.45 | -3.68 | 28 | 934 | 3898 | 2286 |

**Table S2. iKIR score impacts *DQ6* associated protection in a dose-dependent manner.** Individuals were stratified into High, Intermediate and Low iKIR score categories using 6 different definitions of high, intermediate and low (i.e. 6 different strata choices). For all strata choices we observe the same picture: *DQ6* protection increases as iKIR score decreases.

| Haplotype | Group | lnOR | 2.50% | 97.50% | P-value | N haplotype + | | N haplotype - | |
|---|---|---|---|---|---|---|---|---|---|
| | | | | | | Cases | Controls | Cases | Controls |
| ***DRB1*01:01-DQB1*05:01*** | Whole cohort | -0.24 | -0.34 | -0.15 | 3.07E-07 | 1021 | 1150 | 5198 | 4592 |
| | iKIR high | -0.05 | -0.18 | 0.08 | 9.88E-05 | 545 | 644 | 2258 | 2540 |
| | iKIR low | -0.42 | -0.56 | -0.28 | | 476 | 506 | 2940 | 2052 |
| ***DRB1*01:02-DQB1*05:01*** | Whole cohort | -0.34 | -0.65 | -0.03 | 3.18E-02 | 73 | 94 | 6146 | 5648 |
| | iKIR high | -0.01 | -0.47 | 0.45 | 4.20E-02 | 35 | 40 | 2768 | 3144 |
| | iKIR low | -0.65 | -1.08 | -0.24 | | 38 | 54 | 3378 | 2504 |
| ***DRB1*01:03-DQB1*03:01*** | Whole cohort | -1.83 | -2.56 | -1.20 | 9.77E-08 | 10 | 57 | 6209 | 5685 |
| | iKIR high | -1.45 | -2.44 | -0.64 | 2.73E-01 | 6 | 29 | 2797 | 3155 |
| | iKIR low | -2.26 | -3.48 | -1.32 | | 4 | 28 | 3412 | 2530 |
| ***DRB1*04:01-DQB1*03:01*** | Whole cohort | -0.49 | -0.61 | -0.38 | 1.42E-16 | 538 | 772 | 5681 | 4970 |
| | iKIR high | -0.21 | -0.35 | -0.08 | 1.02E-07 | 429 | 583 | 2374 | 2601 |
| | iKIR low | -0.88 | -1.12 | -0.64 | | 109 | 189 | 3307 | 2369 |
| ***DRB1*04:07-DQB1*03:01*** | Whole cohort | -1.95 | -2.47 | -1.49 | 3.22E-15 | 19 | 121 | 6200 | 5621 |
| | iKIR high | -1.59 | -2.26 | -1.01 | 1.26E-01 | 12 | 66 | 2791 | 3118 |
| | iKIR low | -2.38 | -3.26 | -1.66 | | 7 | 55 | 3409 | 2503 |
| ***DRB1*07:01-DQB1*02:02*** | Whole cohort | -1.01 | -1.12 | -0.90 | 2.48E-73 | 542 | 1189 | 5677 | 4553 |
| | iKIR high | -0.87 | -1.01 | -0.73 | 4.47E-02 | 321 | 751 | 2482 | 2433 |
| | iKIR low | -1.09 | -1.27 | -0.92 | | 221 | 438 | 3195 | 2120 |
| ***DRB1*07:01-DQB1*03:03*** | Whole cohort | -2.88 | -3.26 | -2.55 | 2.54E-57 | 33 | 501 | 6186 | 5241 |
| | iKIR high | -2.89 | -3.33 | -2.49 | 2.14E-01 | 24 | 427 | 2779 | 2757 |
| | iKIR low | -2.43 | -3.19 | -1.79 | | 9 | 74 | 3407 | 2484 |
| ***DRB1*10:01-DQB1*05:01*** | Whole cohort | -1.67 | -2.29 | -1.13 | 1.20E-08 | 14 | 69 | 6205 | 5673 |
| | iKIR high | -1.55 | -2.29 | -0.92 | 8.49E-01 | 10 | 53 | 2793 | 3131 |
| | iKIR low | -1.66 | -2.92 | -0.66 | | 4 | 16 | 3412 | 2542 |
| ***DRB1*11:01-DQB1*03:01*** | Whole cohort | -1.77 | -1.99 | -1.55 | 2.38E-57 | 101 | 508 | 6118 | 5234 |
| | iKIR high | -1.70 | -2.00 | -1.42 | 7.02E-01 | 55 | 315 | 2748 | 2869 |
| | iKIR low | -1.78 | -2.12 | -1.47 | | 46 | 193 | 3370 | 2365 |
| ***DRB1*11:04-DQB1*03:01*** | Whole cohort | -1.88 | -2.51 | -1.32 | 4.20E-10 | 13 | 77 | 6206 | 5665 |
| | iKIR high | -1.52 | -2.36 | -0.82 | 2.25E-01 | 8 | 41 | 2795 | 3143 |
| | iKIR low | -2.28 | -3.35 | -1.44 | | 5 | 36 | 3411 | 2522 |
| ***DRB1*12:01-DQB1*03:01*** | Whole cohort | -1.08 | -1.38 | -0.80 | 9.92E-14 | 66 | 176 | 6153 | 5566 |
| | iKIR high | -0.84 | -1.19 | -0.52 | 7.73E-02 | 49 | 126 | 2754 | 3058 |
| | iKIR low | -1.37 | -1.95 | -0.83 | | 17 | 50 | 3399 | 2508 |
| ***DRB1*13:01-DQB1*06:03*** | Whole cohort | -1.49 | -1.68 | -1.31 | 2.34E-56 | 149 | 563 | 6070 | 5179 |

| | | | | | | | | | |
|---|---|---|---|---|---|---|---|---|---|
| | iKIR high | -1.34 | -1.59 | -1.10 | 1.50E-01 | 86 | 342 | 2717 | 2842 |
| | iKIR low | -1.61 | -1.90 | -1.33 | | 63 | 221 | 3353 | 2337 |
| ***DRB1*13:02-DQB1*06:09*** | Whole cohort | -1.50 | -1.91 | -1.12 | 1.04E-13 | 31 | 126 | 6188 | 5616 |
| | iKIR high | -1.50 | -2.20 | -0.89 | 8.52E-01 | 11 | 55 | 2792 | 3129 |
| | iKIR low | -1.57 | -2.10 | -1.09 | | 20 | 71 | 3396 | 2487 |
| ***DRB1*13:03-DQB1*03:01*** | Whole cohort | -2.15 | -2.73 | -1.65 | 5.05E-15 | 15 | 118 | 6204 | 5624 |
| | iKIR high | -2.04 | -2.92 | -1.33 | 6.69E-01 | 7 | 61 | 2796 | 3123 |
| | iKIR low | -2.28 | -3.10 | -1.60 | | 8 | 57 | 3408 | 2501 |
| ***DRB1*14:01-DQB1*05:03*** | Whole cohort | -2.94 | -3.49 | -2.47 | 5.08E-30 | 16 | 265 | 6203 | 5477 |
| | iKIR high | -2.74 | -3.54 | -2.09 | 4.46E-01 | 8 | 134 | 2795 | 3050 |
| | iKIR low | -3.14 | -3.94 | -2.49 | | 8 | 131 | 3408 | 2427 |
| ***DRB1*15:01-DQB1*06:02*** | Whole cohort | -3.75 | -4.04 | -3.48 | 2.11E-155 | 53 | 1533 | 6166 | 4209 |
| | iKIR high | -3.26 | -3.64 | -2.91 | 6.17E-04 | 31 | 717 | 2772 | 2467 |
| | iKIR low | -4.28 | -4.74 | -3.88 | | 22 | 816 | 3394 | 1742 |
| ***DRB1*15:02-DQB1*06:01*** | Whole cohort | -1.71 | -2.42 | -1.11 | 2.24E-07 | 11 | 55 | 6208 | 5687 |
| | iKIR high | -1.41 | -2.31 | -0.65 | 3.33E-01 | 7 | 32 | 2796 | 3152 |
| | iKIR low | -2.08 | -3.31 | -1.12 | | 4 | 23 | 3412 | 2535 |

**Table S1. Impact of functional iKIR on 17 significantly protective *DRB1-DQB1* haplotypes.** iKIR score effect on HLA class II mediated protection was assessed for 17 significantly protective phased haplotypes in our cohort by stratifying the cohort into individuals with a high iKIR score and individuals with a low iKIR score and the protective effect of the haplotype calculated separately in the two strata. Regression coefficients, 95% confidence intervals, p-values and counts in each stratum are reported for each haplotype. The protective effect of class II haplotypes is enhanced in the group of individuals with a low iKIR score (an iKIR score equal to 1.75 or lower) with the exception of *DRB1*07:01-DQB1*03:03*. The odds of seeing this difference by chance were assessed by permutation test for each haplotype ($3x10^7$ permutations). Wald test p-values are reported for the unstratified analysis (Group=Whole cohort).

| | genotype | ln OR | p value | N cases | N controls |
|---|---|---|---|---|---|
| protective | *DQA1*01:02-DQA1*01:03* | -4.24 | 2.63E-09 | 2 | 125 |
| | *DQA1*01:02-DQB1*06:03* | -3.85 | 4.30E-11 | 3 | 128 |
| | *DQA1*01:02-DQB1*06:02* | -3.74 | 1.10E-157 | 54 | 1545 |
| | *DQA1*01:01-DQA1*05:05-DQB1*03:01-DQB1*05:01* | -3.08 | 2.00E-13 | 6 | 118 |
| | *DRB1*14:01-DQA1*01:04-DQB1*05:03* | -2.94 | 5.10E-30 | 16 | 265 |
| | *DQA1*02:01-DQB1*03:03* | -2.60 | 3.10E-63 | 46 | 524 |
| | *DQA1*01:02-DQB1*05:01* | -2.08 | 5.20E-37 | 43 | 304 |
| | *DRB1*07:01-DQA1*02:01-DQB1*05:01* | -2.06 | 3.90E-28 | 33 | 232 |
| | *DRB1*04:07-DQA1*03:03-DQB1*03:01* | -1.95 | 3.20E-15 | 19 | 121 |
| | *DQA1*01:02* | -1.80 | 5.60E-249 | 526 | 2051 |
| | *DQA1*05:05-DQB1*03:01* | -1.70 | 9.20E-104 | 213 | 933 |
| | *DRB1*10:01-DQA1*01:05-DQB1*05:01* | -1.67 | 1.20E-08 | 14 | 69 |
| | *B*57:01* | -1.67 | 5.90E-52 | 103 | 474 |
| | *DQA1*01:03* | -1.51 | 8.80E-63 | 161 | 618 |
| | *DQA1*02:01* | -1.37 | 8.50E-149 | 573 | 1639 |
| | *DQB1*03:01* | -1.16 | 4.10E-129 | 759 | 1773 |
| | *C*08:02* | -0.94 | 4.00E-29 | 215 | 484 |
| | *A*26:01* | -0.63 | 9.50E-09 | 137 | 232 |
| | *C*04:01* | -0.54 | 9.20E-22 | 593 | 882 |
| | *A*32:01* | -0.53 | 1.30E-11 | 283 | 432 |
| | *A*11:01* | -0.50 | 9.60E-16 | 472 | 687 |
| | **genotype** | **ln OR** | **p value** | **N cases** | **N controls** |
| detrimental | *A*24:02* | 0.45 | 1.10E-19 | 1284 | 820 |
| | *B*18:01* | 0.73 | 9.20E-30 | 827 | 397 |
| | *A*30:02* | 1.08 | 3.20E-20 | 305 | 98 |
| | *DRB1*04:01-DQA1*01:01* | 1.25 | 2.10E-39 | 522 | 146 |
| | *DRB1*03:01-DQA1*05:01-DQB1*02:01* | 1.34 | 7.10E-257 | 3724 | 1612 |
| | *B*39:06* | 1.48 | 1.10E-31 | 356 | 78 |
| | *DRB1*04:05* | 1.86 | 7.90E-28 | 270 | 40 |
| | *DRB1*04:01-DQB1*03:02* | 2.01 | 0.00E+00 | 2849 | 583 |
| | *DQB1*03:02* | 2.04 | 0.00E+00 | 3996 | 1085 |
| | *DRB1*03:01-DQA1*05:01-DQB1*02:01-DQB1*03:02* | 2.94 | 7.20E-264 | 2177 | 159 |

**Table S2. The HLA class I and class II genotypes most closely identified with outcome in the GRID cohort.** This table shows the “driver” genotypes which were independently associated with outcome (see **Fig. 3**). The natural log of the odds ratio (ln[OR]), associated p-value for each genotype and number of cases and controls carrying the genotype is shown. ln[OR] and p-values were obtained by multiple logistic regression in the whole cohort with gender as an additional covariate is reported (i.e. the coefficients and p-values are derived in a model containing one genotype at a time).

| Protective genotype | Coefficient of interaction | P-value of interaction | N cases | N controls | N total |
|---|---|---|---|---|---|
| *DQA1*01:02-DQB1*06:02* | 0.94 | 1.14E-05 | 26 | 729 | 755 |
| *DQB1*03:01* | 0.35 | 1.97E-05 | 361 | 777 | 1138 |
| *DQA1*01:02* | 0.33 | 8.35E-05 | 252 | 922 | 1174 |
| *DQA1*05:05-DQB1*03:01* | 0.30 | 3.53E-02 | 88 | 355 | 443 |
| *DQA1*01:02-DQB1*05:01* | 0.65 | 5.54E-02 | 256 | 548 | 804 |
| *DQA1*02:01* | 0.18 | 7.35E-02 | 14 | 109 | 123 |
| *DQA1*01:03* | 0.24 | 1.19E-01 | 67 | 279 | 346 |
| *DRB1*07:01-DQA1*02:01-DQB1*05:01* | 0.60 | 2.27E-01 | 12 | 52 | 64 |
| *DQA1*02:01-DQB1*03:03* | 0.11 | 7.31E-01 | 12 | 85 | 97 |

**Table S3. iKIR score interaction terms with protective HLA class II drivers in a subcohort without HLA class I drivers.** After removal of class I drivers from the cohort (remaining cohort: N=5,420), we modelled risk of T1D for each protective genotype and included iKIR score in the model as a continuous variable interacting with the HLA class II genotype ($\boldsymbol{OUTCOME} \sim \boldsymbol{HLA\ class\ II\ genotype} \times \boldsymbol{iKIR\_score} + \boldsymbol{GENDER}$). The coefficient for the interaction term was in the expected direction for all genotypes and significant for 4 frequent genotypes.

| Haplotype | Threshold | Group | lnOR | 2.50% | 97.50% | P-value | N Genotype + | | N Genotype - | |
|---|---|---|---|---|---|---|---|---|---|---|
| | | | | | | | Cases | Controls | Cases | Controls |
| ***DQA1*01:02*** | Unstratified | Whole cohort | -1.91 | -2.07 | -1.76 | $8.14x10^{-133}$ | 252 | 922 | 2758 | 1488 |
| | 1.5 | iKIR high | -1.75 | -1.94 | -1.56 | $5.94\ x10^{-5}$ | 155 | 619 | 1725 | 1202 |
| | | iKIR low | -2.42 | -2.69 | -2.16 | | 97 | 303 | 1033 | 286 |
| | 1.75 | iKIR high | -1.63 | -1.88 | -1.38 | $7.14x10^{-4}$ | 88 | 359 | 999 | 802 |
| | | iKIR low | -2.18 | -2.38 | -1.98 | | 164 | 563 | 1759 | 686 |
| | 2 | iKIR high | -1.58 | -1.85 | -1.33 | $5.13x10^{-4}$ | 82 | 318 | 887 | 706 |
| | | iKIR low | -2.14 | -2.33 | -1.95 | | 170 | 604 | 1871 | 782 |
| ***DQA1*01:02-DQB1*05:01*** | Unstratified | Whole cohort | -2.32 | -2.92 | -1.80 | $4.14x10^{-16}$ | 14 | 109 | 2996 | 2301 |
| | 1.5 | iKIR high | -2.00 | -2.70 | -1.41 | 0.12 | 11 | 76 | 1869 | 1745 |
| | | iKIR low | -3.10 | -4.53 | -2.07 | | 3 | 33 | 1127 | 556 |
| | 1.75 | iKIR high | -1.80 | -2.69 | -1.06 | 0.14 | 7 | 44 | 1080 | 1117 |
| | | iKIR low | -2.71 | -3.58 | -1.99 | | 7 | 65 | 1916 | 1184 |
| | 2 | iKIR high | -1.74 | -2.64 | -1.00 | 0.13 | 7 | 41 | 962 | 983 |
| | | iKIR low | -2.71 | -3.58 | -1.99 | | 7 | 68 | 2034 | 1318 |
| ***DQA1*01:02-DQB1*06:02*** | Unstratified | Whole cohort | -3.91 | -4.33 | -3.53 | $1.76x10^{-83}$ | 26 | 729 | 2984 | 1681 |
| | 1.5 | iKIR high | -3.45 | -3.92 | -3.03 | $3.64x10^{-3}$ | 21 | 477 | 1859 | 1344 |
| | | iKIR low | -5.12 | -6.16 | -4.33 | | 5 | 252 | 1125 | 337 |
| | 1.75 | iKIR high | -3.12 | -3.69 | -2.63 | $1.81x10^{-3}$ | 15 | 279 | 1072 | 882 |
| | | iKIR low | -4.58 | -5.25 | -4.03 | | 11 | 450 | 1912 | 799 |
| | 2 | iKIR high | -3.09 | -3.68 | -2.58 | $4.06x10^{-3}$ | 14 | 249 | 955 | 775 |
| | | iKIR low | -4.49 | -5.13 | -3.96 | | 12 | 480 | 2029 | 906 |
| ***DQA1*01:03*** | Unstratified | Whole cohort | -1.75 | -2.03 | -1.49 | $2.12x10^{-36}$ | 67 | 279 | 2943 | 2131 |
| | 1.5 | iKIR high | -1.58 | -1.90 | -1.28 | 0.061 | 52 | 221 | 1828 | 1600 |
| | | iKIR low | -2.08 | -2.70 | -1.53 | | 15 | 58 | 1115 | 531 |
| | 1.75 | iKIR high | -1.54 | -1.93 | -1.18 | 0.26 | 35 | 156 | 1052 | 1005 |
| | | iKIR low | -1.86 | -2.27 | -1.48 | | 32 | 123 | 1891 | 1126 |
| | 2 | iKIR high | -1.38 | -1.78 | -1.01 | 0.036 | 35 | 133 | 934 | 891 |
| | | iKIR low | -2.00 | -2.41 | -1.63 | | 32 | 146 | 2009 | 1240 |
| ***DQA1*02:01*** | Unstratified | Whole cohort | -1.15 | -1.31 | -0.99 | $2.02x10^{-45}$ | 256 | 548 | 2754 | 1862 |
| | 1.5 | iKIR high | -0.97 | -1.15 | -0.80 | $5.14x10^{-3}$ | 235 | 500 | 1645 | 1321 |
| | | iKIR low | -1.54 | -2.08 | -1.03 | | 21 | 48 | 1109 | 541 |
| | 1.75 | iKIR high | -0.88 | -1.10 | -0.67 | $6.15x10^{-3}$ | 149 | 322 | 938 | 839 |
| | | iKIR low | -1.32 | -1.57 | -1.08 | | 107 | 226 | 1816 | 1023 |
| | 2 | iKIR high | -0.93 | -1.16 | -0.70 | 0.044 | 130 | 288 | 839 | 736 |

| | | | | | | | | | | |
|---|---|---|---|---|---|---|---|---|---|---|
| | | iKIR low | -1.25 | -1.48 | -1.03 | | 126 | 260 | 1915 | 1126 |
| ***DQA1*02:01-DQB1*03:03*** | Unstratified | Whole cohort | -2.21 | -2.87 | -1.65 | $9.05 \times 10^{-13}$ | 12 | 85 | 2998 | 2325 |
| | 1.5 | iKIR high | -2.12 | -2.89 | -1.48 | 0.92 | 9 | 70 | 1871 | 1751 |
| | | iKIR low | -2.26 | -3.73 | -1.15 | | 3 | 15 | 1127 | 574 |
| | 1.75 | iKIR high | -2.25 | -3.31 | -1.43 | 0.8 | 5 | 49 | 1082 | 1112 |
| | | iKIR low | -2.09 | -2.99 | -1.34 | | 7 | 36 | 1916 | 1213 |
| | 2 | iKIR high | -2.11 | -3.18 | -1.27 | 0.86 | 5 | 42 | 964 | 982 |
| | | iKIR low | -2.23 | -3.12 | -1.49 | | 7 | 43 | 2034 | 1343 |
| ***DQA1*05:05-DQB1*03:01*** | Unstratified | Whole cohort | -1.74 | -1.99 | -1.51 | $6.01 \times 10^{-46}$ | 88 | 355 | 2922 | 2055 |
| | 1.5 | iKIR high | -1.63 | -1.90 | -1.37 | $2.96 \times 10^{-3}$ | 70 | 300 | 1810 | 1521 |
| | | iKIR low | -1.85 | -2.42 | -1.33 | | 18 | 55 | 1112 | 534 |
| | 1.75 | iKIR high | -1.39 | -1.72 | -1.08 | $7.49 \times 10^{-3}$ | 53 | 198 | 1034 | 963 |
| | | iKIR low | -2.05 | -2.44 | -1.69 | | 35 | 157 | 1888 | 1092 |
| | 2 | iKIR high | -1.40 | -1.75 | -1.07 | $1.90 \times 10^{-2}$ | 46 | 171 | 923 | 853 |
| | | iKIR low | -1.99 | -2.34 | -1.65 | | 42 | 184 | 1999 | 1202 |
| ***DQB1*03:01*** | Unstratified | Whole cohort | -1.25 | -1.39 | -1.11 | $5.59 \times 10^{-69}$ | 361 | 777 | 2649 | 1633 |
| | 1.5 | iKIR high | -1.05 | -1.20 | -0.90 | $5.09 \times 10^{-6}$ | 319 | 671 | 1561 | 1150 |
| | | iKIR low | -1.73 | -2.12 | -1.37 | | 42 | 106 | 1088 | 483 |
| | 1.75 | iKIR high | -0.85 | -1.03 | -0.67 | $<1 \times 10^{-8}$ | 269 | 505 | 818 | 656 |
| | | iKIR low | -1.71 | -1.96 | -1.46 | | 92 | 272 | 1831 | 977 |
| | 2 | iKIR high | -0.95 | -1.15 | -0.75 | $7.09 \times 10^{-4}$ | 213 | 431 | 756 | 593 |
| | | iKIR low | -1.44 | -1.65 | -1.24 | | 148 | 346 | 1893 | 1040 |
| ***DRB1*07:01-DQA1*02:01-DQB1*05:01*** | Unstratified | Whole cohort | -1.69 | -2.37 | -1.10 | $1.40 \times 10^{-7}$ | 12 | 52 | 2998 | 2358 |
| | 1.5 | iKIR high | -1.48 | -2.16 | -0.88 | $1.14 \times 10^{-2}$ | 12 | 50 | 1868 | 1771 |
| | | iKIR low | -14.22 | - | 45.10 | | 0 | 2 | 1130 | 587 |
| | 1.75 | iKIR high | -1.13 | -1.94 | -0.41 | $6.51 \times 10^{-2}$ | 9 | 29 | 1078 | 1132 |
| | | iKIR low | -2.45 | -3.89 | -1.39 | | 3 | 23 | 1920 | 1226 |
| | 2 | iKIR high | -1.11 | -1.97 | -0.35 | $1.05 \times 10^{-1}$ | 8 | 25 | 961 | 999 |
| | | iKIR low | -2.29 | -3.51 | -1.34 | | 4 | 27 | 2037 | 1359 |

**Table S4. iKIR score decreases protection associated with protective class II genotypes in T1D.** The GRID cohort without carriers of HLA class I drivers (N=5,420) was stratified into individuals with high or low iKIR score at different cutoffs (1.5, 1.75 and 2.0). The protective effect of each protective genotype was evaluated independently in each stratum using multivariate logistic regression with gender as covariate. HLA class II protection is enhanced in the iKIR low strata for all genotypes but for the very infrequent protective genotype *DQA1*02:01-DQB1*03:03*. Regression coefficients, permutation p-values and cohort sizes are reported for the different strata. P-value for the whole cohort (unstratified analysis) calculated using the Wald-test; p-values for the stratification analysis are calculated using the permutation test.

| iKIR score threshold | P-value |
|---|---|
| 1 | $2.9x10^{-03}$ |
| 1.5 | $8.8x10^{-03}$ |
| 1.75 | $1x10^{-05}$ |
| 2 | $2.1x10^{-04}$ |
| 2.5 | $2.2x10^{-03}$ |

**Table S5. Odds of observing an iKIR modification across *DQA1*01:02*, *DQB1*03:01*, *DQA1*02:01* and *DQA1*01:02-DQB1*06:02* in an independent cohort.** The iKIR effect observed in the case-control cohort was validated in an independent family dataset. Trios were stratified into high (>threshold) and low (≤ threshold) iKIR score according to the iKIR score of the child in each trio. For each threshold and each genotype, we calculated the ratio of transmitted to non-transmitted genes in each stratum. The odds of observing an equal or greater difference between log ratios across 4 frequent protective genotypes (*DQA1*01:02*, *DQB1*03:01*, *DQA1*02:01* and *DQA1*01:02-DQB1*06:02*) were assessed by permutation test.

| | Group | ln[OR] | 2.50% | 97.50% | P-value | N haplotype + | | N haplotype- | |
|---|---|---|---|---|---|---|---|---|---|
| | | | | | | Cases | Controls | Cases | Controls |
| | Whole cohort | -3.87 | -4.16 | -3.6 | 2.79E-163 | 53 | 1533 | 6166 | 4209 |
| Threshold=1.5 | iKIR high | -3.56 | -3.88 | -3.28 | 2.29E-03 | 46 | 1156 | 4693 | 3582 |
| | iKIR low | -4.89 | -5.75 | -4.22 | | 7 | 377 | 1473 | 627 |
| Threshold=1.75 | iKIR high | -3.29 | -3.68 | -2.95 | 6.17E-04 | 31 | 717 | 2772 | 2467 |
| | iKIR low | -4.36 | -4.82 | -3.96 | | 22 | 816 | 3394 | 1742 |
| Threshold=2 | iKIR high | -3.33 | -3.74 | -2.96 | 5.45E-03 | 27 | 625 | 2401 | 2071 |
| | iKIR low | -4.24 | -4.66 | -3.86 | | 26 | 908 | 3765 | 2138 |
| Threshold=2.5 | iKIR high | -3.38 | -3.82 | -3.00 | 2.40E-02 | 25 | 607 | 2268 | 1915 |
| | iKIR low | -4.17 | -4.57 | -3.81 | | 28 | 926 | 3898 | 2294 |

**Table S6. *DRB1*15:01-DQB1*06:02* protection in T1D is enhanced amongst individuals with a low iKIR score.** The cohort was stratified into individuals with high or low iKIR score using different iKIR score thresholds (1.5, 1.75, 2.0 and 2.5). The protective effect of *DRB1*15:01-DQB1*06:02* was evaluated independently in each stratum using multivariate logistic regression with gender included as a covariate. *DRB1*15:01-DQB1*06:02* is significantly protective (ln[OR]= -3.9, $P=1.14 \times 10^{-165}$) in the cohort (Group=Whole cohort, unstratified analysis). Regression coefficients, p-values and cohort sizes are reported for the different strata. P-value for the unstratified analysis calculated using Wald-test; all other p-values calculated using a permutation test.

| Strata choice | Group | | ln[OR] | 2.50% | 97.50% | N genotype+ | | N genotype- | |
|---|---|---|---|---|---|---|---|---|---|
| | | | | | | Cases | Controls | Cases | Controls |
| 1 | High | (2.75,4] | -2.32 | -2.95 | -1.78 | 13 | 125 | 514 | 481 |
| | Int | (1.75,2.75] | -3.62 | -4.13 | -3.18 | 18 | 592 | 2258 | 1986 |
| | Low | [0,1.75] | -4.28 | -4.74 | -3.88 | 22 | 816 | 3394 | 1742 |
| 2 | High | (3,4] | -2.34 | -3.00 | -1.77 | 12 | 119 | 470 | 447 |
| | Int | (1.75,3] | -3.58 | -4.08 | -3.15 | 19 | 598 | 2302 | 2020 |
| | Low | [0,1.75] | -4.28 | -4.74 | -3.88 | 22 | 816 | 3394 | 1742 |
| 3 | High | (2.75,4] | -2.32 | -2.95 | -1.78 | 13 | 125 | 514 | 481 |
| | Int | (2,2.75] | -3.75 | -4.33 | -3.25 | 14 | 500 | 1887 | 1590 |
| | Low | [0,2] | -4.12 | -4.54 | -3.75 | 26 | 908 | 3765 | 2138 |
| 4 | High | (3,4] | -2.34 | -3.00 | -1.77 | 12 | 119 | 470 | 447 |
| | Int | (2,3] | -3.70 | -4.26 | -3.21 | 15 | 506 | 1931 | 1624 |
| | Low | [0,2] | -4.12 | -4.54 | -3.75 | 26 | 908 | 3765 | 2138 |
| 5 | High | (2.75,4] | -2.32 | -2.95 | -1.78 | 13 | 125 | 514 | 481 |
| | Int | (2.5,2.75] | -3.90 | -4.53 | -3.37 | 12 | 482 | 1754 | 1434 |
| | Low | [0,2.5] | -4.03 | -4.43 | -3.67 | 28 | 926 | 3898 | 2294 |
| 6 | High | (3,4] | -2.34 | -3.00 | -1.77 | 12 | 119 | 470 | 447 |
| | Int | (2.5,3] | -3.83 | -4.44 | -3.32 | 13 | 488 | 1798 | 1468 |
| | Low | [0,2.5] | -4.03 | -4.43 | -3.67 | 28 | 926 | 3898 | 2294 |

**Table S7. iKIR score impacts *DRB1*15:01-DQB1*06:02* protection in a dose-dependent manner.** Individuals were stratified into High, Intermediate and Low iKIR score categories using 6 different definitions of high, intermediate, and low (i.e. 6 different strata choices). For all strata choices we observe the same picture, that *DRB1*15:01-DQB1*06:02* protection increases as iKIR score decreases as we observed for *DQA1*01:02-DQB1*06:02* (see **Fig. S3**).

| Covariates | Coefficient of interaction | P-value of interaction |
|---|---|---|
| GENDER + *DR3* | +0.67 | $1.9x10^{-6}$ |
| GENDER + *DR4* | +0.70 | $1x10^{-6}$ |
| GENDER + *DR3* + *DR4* | +0.67 | $3.4x10^{-6}$ |

**Table S8. iKIR score effect on *DR*15:01-DQ*06:02* is independent of *DR3* and *DR4* haplotypes.** *DR3* and *DR4* haplotypes were included as covariates and standardised iKIR score was included as an interaction term for comparison. We denote the strata *DR3* or *DR4* for ease of reference, but we are considering genes both in cis or in trans and only considering *DRB1* and *DQB1* genes.

| Allele | ln[OR] | P-value | Allele | ln[OR] | P-value | Allele | ln[OR] | P-value | Allele | ln[OR] | P-value |
|---|---|---|---|---|---|---|---|---|---|---|---|
| *A0101* | 0.54 | 2.80E-03 | *B0801* | 0.66 | 2.16E-04 | *B5501* | 0.74 | 4.11E-06 | *C0304* | 0.77 | 1.42E-05 |
| *A0201* | 0.77 | 2.31E-03 | *B1501* | 0.72 | 3.10E-05 | *B4901* | 0.79 | 9.39E-07 | *C0303* | 0.79 | 2.45E-06 |
| *A2402* | 0.86 | 3.21E-06 | *B4402* | 0.81 | 2.64E-06 | *B4002* | 0.76 | 1.62E-06 | *C0501* | 0.88 | 3.79E-07 |
| *A3201* | 0.79 | 9.36E-07 | *B3501* | 0.73 | 5.55E-06 | *B4101* | 0.75 | 2.08E-06 | *C0701* | 0.70 | 1.48E-04 |
| *A1101* | 0.76 | 3.62E-06 | *B1401* | 0.79 | 1.12E-06 | *B1503* | 0.75 | 2.35E-06 | *C0401* | 0.74 | 3.46E-06 |
| *A2301* | 0.75 | 2.16E-06 | *B4403* | 0.74 | 4.64E-06 | *B2702* | 0.75 | 2.29E-06 | *C0802* | 0.84 | 3.86E-07 |
| *A0301* | 0.92 | 6.42E-05 | *B0702* | 1.00 | 6.02E-04 | *B4501* | 0.75 | 2.22E-06 | *C0602* | 0.61 | 3.03E-04 |
| *A2902* | 0.73 | 6.21E-06 | *B3901* | 0.74 | 2.93E-06 | *B4102* | 0.75 | 2.56E-06 | *C1203* | 0.71 | 1.52E-05 |
| *A2501* | 0.72 | 1.33E-05 | *B1801* | 0.76 | 6.65E-06 | *B4405* | 0.75 | 2.09E-06 | *C0202* | 0.74 | 5.35E-06 |
| *A0205* | 0.77 | 1.94E-06 | *B3801* | 0.75 | 2.44E-06 | *B1510* | 0.75 | 2.29E-06 | *C0702* | 1.17 | 5.29E-04 |
| *A3002* | 0.76 | 1.73E-06 | *B4001* | 0.84 | 9.62E-07 | *B5701* | 0.67 | 2.91E-05 | *C0302* | 0.75 | 2.29E-06 |
| *A0206* | 0.75 | 2.23E-06 | *B3906* | 0.78 | 4.48E-06 | *B5301* | 0.75 | 2.33E-06 | *C0102* | 0.72 | 6.13E-06 |
| *A2601* | 0.74 | 2.73E-06 | *B1302* | 0.77 | 2.16E-06 | *B5108* | 0.75 | 2.31E-06 | *C1601* | 0.72 | 6.84E-06 |
| *A3101* | 0.77 | 4.44E-06 | *B2705* | 0.71 | 1.15E-05 | *B1516* | 0.75 | 2.29E-06 | *C0704* | 0.75 | 2.40E-06 |
| *A3001* | 0.74 | 3.43E-06 | *B1402* | 0.80 | 8.50E-07 | *B1508* | 0.75 | 2.30E-06 | *C1502* | 0.71 | 1.26E-05 |
| *A6801* | 0.74 | 3.01E-06 | *B5201* | 0.72 | 7.70E-06 | *B4006* | 0.75 | 2.26E-06 | *C1202* | 0.72 | 7.72E-06 |
| *A0202* | 0.75 | 2.25E-06 | *B5101* | 0.72 | 8.95E-06 | *B4801* | 0.75 | 2.29E-06 | *C1402* | 0.74 | 2.71E-06 |
| *A3301* | 0.75 | 2.20E-06 | *B1517* | 0.74 | 3.66E-06 | *B5601* | 0.75 | 2.25E-06 | *C1505* | 0.75 | 2.13E-06 |
| *A3004* | 0.75 | 2.33E-06 | *B4701* | 0.77 | 1.47E-06 | *B3924* | 0.75 | 2.24E-06 | *C0210* | 0.75 | 2.29E-06 |
| *A2901* | 0.75 | 2.38E-06 | *B5001* | 0.76 | 1.67E-06 | *B5801* | 0.75 | 2.20E-06 | *C1701* | 0.75 | 2.40E-06 |
| *A6601* | 0.75 | 2.25E-06 | *B3701* | 0.66 | 4.66E-05 | *B7301* | 0.75 | 2.26E-06 | *C1602* | 0.75 | 2.06E-06 |
| *A3303* | 0.75 | 2.41E-06 | *B3503* | 0.75 | 2.24E-06 | | | | *C1604* | 0.75 | 2.24E-06 |
| *A3402* | 0.75 | 2.34E-06 | *B1518* | 0.75 | 2.57E-06 | | | | *C1403* | 0.75 | 2.29E-06 |
| *A6802* | 0.75 | 2.13E-06 | *B0705* | 0.75 | 2.17E-06 | | | | *C0310* | 0.75 | 2.32E-06 |
| *A6901* | 0.75 | 2.30E-06 | *B3502* | 0.75 | 2.04E-06 | | | | *C0803* | 0.75 | 2.29E-06 |
| *A7403* | 0.75 | 2.29E-06 | *B3508* | 0.76 | 1.92E-06 | | | | | | |

**Table S9. iKIR interaction remains significant in all HLA class I allele negative subcohorts**. Individuals carrying a given HLA class I allele (Allele) are removed from the GRID cohort and then the subcohort is modeled with iKIR as an interaction term with *DRB1*15:01-DQB1*06:02*. Coefficients (ln[OR]) and p-values for the interaction term are reported.

| | Group | ln[OR] | 2.50% | 97.50% | P-value | N haplotype + | | N haplotype- | |
|---|---|---|---|---|---|---|---|---|---|
| | | | | | | Cases | Controls | Cases | Controls |
| | Whole cohort | -3.86 | -4.15 | -3.59 | $1.3 \times 10^{-165}$ | 54 | 1545 | 6165 | 4197 |
| Threshold=1.5 | iKIR high | -3.55 | -3.86 | -3.27 | $2.0 \times 10^{-3}$ | 47 | 1165 | 4692 | 3573 |
| | iKIR low | -4.27 | -4.80 | -3.82 | | 7 | 380 | 1473 | 624 |
| Threshold=1.75 | iKIR high | -3.27 | -3.65 | -2.93 | $2.8 \times 10^{-4}$ | 32 | 721 | 2771 | 2463 |
| | iKIR low | -4.38 | -4.84 | -3.98 | | 22 | 824 | 3394 | 1734 |
| Threshold=2 | iKIR high | -3.30 | -3.70 | -2.93 | $2.6 \times 10^{-3}$ | 28 | 629 | 2400 | 2067 |
| | iKIR low | -4.25 | -4.67 | -3.88 | | 26 | 916 | 3765 | 2130 |
| Threshold=2.5 | iKIR high | -3.35 | -3.78 | -2.98 | $9.2 \times 10^{-3}$ | 26 | 611 | 2267 | 1911 |
| | iKIR low | -4.18 | -4.59 | -3.82 | | 28 | 934 | 3898 | 2286 |

**Table S10. iKIR score negatively impacts protection associated with *DQ6* in T1D even when iKIR ligands are included as covariates.** The GRID cohort was stratified into individuals with high or low iKIR score using different cutoffs (1.5, 1.75, 2.0 and 2.5). The protective effect of *DQ6* was evaluated independently in each stratum using multivariate logistic regression with gender, Bw4, C1 and C2 ligands included in the model as covariates. Overall conclusions were remarkably similar to our previous analysis (not including the ligands as covariates). Regression coefficients, permutation p-values and cohort sizes are reported for the different strata. P-value for the whole cohort (unstratified analysis) calculated using the Wald-test; p-values for the stratification analysis are calculated using the permutation test.

| Allele | ln[OR] | P-value | Allele | ln[OR] | P-value | Allele | ln[OR] | P-value | Allele | ln[OR] | P-value |
|---|---|---|---|---|---|---|---|---|---|---|---|
| *A0101* | 0.54 | 2.86E-03 | *B0801* | 0.66 | 2.24E-04 | *B5501* | 0.77 | 1.19E-06 | *C0304* | 0.81 | 4.24E-06 |
| *A0201* | 0.86 | 5.78E-04 | *B1501* | 0.75 | 9.79E-06 | *B4901* | 0.82 | 2.60E-07 | *C0303* | 0.83 | 6.72E-07 |
| *A2402* | 0.91 | 6.95E-07 | *B4402* | 0.85 | 6.18E-07 | *B4002* | 0.80 | 4.57E-07 | *C0501* | 0.92 | 7.82E-08 |
| *A3201* | 0.83 | 2.54E-07 | *B3501* | 0.76 | 1.61E-06 | *B4101* | 0.79 | 5.92E-07 | *C0701* | 0.70 | 1.53E-04 |
| *A1101* | 0.79 | 1.09E-06 | *B1401* | 0.82 | 3.09E-07 | *B1503* | 0.78 | 6.75E-07 | *C0401* | 0.78 | 9.90E-07 |
| *A2301* | 0.78 | 6.19E-07 | *B4403* | 0.77 | 1.39E-06 | *B2702* | 0.79 | 6.55E-07 | *C0802* | 0.88 | 1.06E-07 |
| *A0301* | 0.98 | 1.81E-05 | *B0702* | 1.09 | 1.47E-04 | *B4501* | 0.78 | 6.35E-07 | *C0602* | 0.66 | 8.85E-05 |
| *A2902* | 0.76 | 1.86E-06 | *B3901* | 0.78 | 8.48E-07 | *B4102* | 0.78 | 7.39E-07 | *C1203* | 0.75 | 4.46E-06 |
| *A2501* | 0.76 | 3.87E-06 | *B1801* | 0.80 | 1.89E-06 | *B4405* | 0.79 | 5.97E-07 | *C0202* | 0.78 | 1.48E-06 |
| *A0205* | 0.81 | 5.45E-07 | *B3801* | 0.78 | 7.05E-07 | *B1510* | 0.79 | 6.56E-07 | *C0702* | 1.28 | 1.16E-04 |
| *A3002* | 0.79 | 4.89E-07 | *B4001* | 0.87 | 2.57E-07 | *B5701* | 0.71 | 8.42E-06 | *C0302* | 0.79 | 6.55E-07 |
| *A0206* | 0.79 | 6.37E-07 | *B3906* | 0.82 | 1.18E-06 | *B5301* | 0.78 | 6.68E-07 | *C0102* | 0.76 | 1.83E-06 |
| *A2601* | 0.78 | 7.90E-07 | *B1302* | 0.81 | 5.98E-07 | *B5108* | 0.78 | 6.63E-07 | *C1601* | 0.76 | 2.07E-06 |
| *A3101* | 0.77 | 4.09E-06 | *B2705* | 0.75 | 3.33E-06 | *B1516* | 0.78 | 6.57E-07 | *C0704* | 0.78 | 7.03E-07 |
| *A3001* | 0.78 | 9.89E-07 | *B1402* | 0.83 | 2.43E-07 | *B1508* | 0.79 | 6.59E-07 | *C1502* | 0.71 | 1.17E-05 |
| *A6801* | 0.78 | 8.75E-07 | *B5201* | 0.75 | 2.28E-06 | *B4006* | 0.79 | 6.45E-07 | *C1202* | 0.75 | 2.28E-06 |
| *A0202* | 0.79 | 6.43E-07 | *B5101* | 0.72 | 8.36E-06 | *B4801* | 0.79 | 6.55E-07 | *C1402* | 0.78 | 7.86E-07 |
| *A3301* | 0.79 | 6.27E-07 | *B1517* | 0.78 | 1.05E-06 | *B5601* | 0.79 | 6.43E-07 | *C1505* | 0.79 | 6.08E-07 |
| *A3004* | 0.78 | 6.68E-07 | *B4701* | 0.80 | 4.11E-07 | *B3924* | 0.79 | 6.41E-07 | *C0210* | 0.79 | 6.56E-07 |
| *A2901* | 0.78 | 6.83E-07 | *B5001* | 0.79 | 4.71E-07 | *B5801* | 0.79 | 6.30E-07 | *C1701* | 0.78 | 6.90E-07 |
| *A6601* | 0.79 | 6.45E-07 | *B3701* | 0.70 | 1.43E-05 | *B7301* | 0.79 | 6.46E-07 | *C1602* | 0.79 | 5.86E-07 |
| *A3303* | 0.78 | 6.90E-07 | *B3503* | 0.78 | 6.42E-07 | | | | *C1604* | 0.79 | 6.42E-07 |
| *A3402* | 0.78 | 6.69E-07 | *B1518* | 0.78 | 7.50E-07 | | | | *C1403* | 0.79 | 6.56E-07 |
| *A6802* | 0.79 | 6.06E-07 | *B0705* | 0.79 | 6.20E-07 | | | | *C0310* | 0.78 | 6.66E-07 |
| *A6901* | 0.79 | 6.59E-07 | *B3502* | 0.79 | 5.81E-07 | | | | *C0803* | 0.79 | 6.55E-07 |
| *A7403* | 0.79 | 6.55E-07 | *B3508* | 0.80 | 5.36E-07 | | | | | | |

**Table S11. iKIR interaction remains significant in all HLA class I allele negative subcohorts**. Individuals carrying a given HLA class I allele (Allele) are removed from the GRID cohort and then the risk of T1D is modeled with iKIR as an interaction term with *DQ6* in the allele-negative subcohort. Coefficients (ln[OR]) and p-values for the interaction term are reported.

| Covariates | Coefficient of interaction | P-value of interaction |
|---|---|---|
| GENDER + *DR3* | +0.69 | $5.3 \times 10^{-7}$ |
| GENDER + *DR4* | +0.73 | $3.4 \times 10^{-7}$ |
| GENDER + *DR3* + *DR4* | +0.69 | $1.3 \text{X} 10^{-6}$ |

**Table S12. iKIR score effect on *DQ6* is independent of *DR3* and *DR4* detrimental genotypes.** *DR3* (defined here to be *DRB1*03:01-DQB1*02:01 in cis* or *in trans*) and *DR4* (defined to be *DRB1*04:01/02/04/05-DQB1*03:02 in cis* or *in trans*) detrimental genotypes were included as covariates and standardised iKIR score was included as an interaction term for comparison.

| Equation | Parameter | Model | Definition | Units | Value | Reference |
|---|---|---|---|---|---|---|
| β-cell (B) | $\delta_\beta$ | 1&2 | β cell killing per effector T cell | $cell^{-1}day^{-1}$ | $1x10^{-5}$ – $9x10^{-5}$ | Estimated |
| Islet antigen (A) | $\alpha_A$ | 1&2 | Rate of islet antigen release (or beta cell damage) per Tconv | $cell^{-1}day^{-1}$ | $1x10^{-5}$ | 33 |
| | $\delta_A$ | 1&2 | Clearance rate of the antigen | $day^{-1}$ | 1 | 33 |
| $T_{regs}$ (R) | $\alpha_R$ | 1&2 | Proliferation rate of Tregs upon antigen encounter | $day^{-1}$/ $cell^{-1}day^{-1}$ | 1 – 10 | 34 |
| | $\delta_T$ | 1&2 | Death rate of Tregs and Tconvs | $day^{-1}$ | 0.1 – 0.2 | 35 |
| | $K_R$ | 2 | Density control on Treg proliferation rate | cells | $\gamma_R$ – 10 | 34 |
| $T_{convs}$ (C) | $\alpha_C$ | 1&2 | Proliferation rate of Tconvs dependent of antigen activation of Tnaive and memory cells | $day^{-1}$ | 1 – 100 | 34 |
| | $\delta_i$ | 1 | T conv suppression by Tregs | $cell^{-1}day^{-1}$ | 1 | Estimated |
| | k | 2 | Threshold of Tconv inhibition by Treg | cells | 1 | Estimated |
| | $K_C$ | 2 | Density control on Tconv proliferation rate | cells | $\gamma_C$ – 100 | 34 |

**Table S13. Parameters used in the mathematical model of Beta cell destruction.**

| Population | Antibody | Supplier | Product code |
|---|---|---|---|
| Dump | CD14 V500 | BD Horizon | 561391 |
| | CD19 V500 | BD Horizon | 561121 |
| | Live/dead fixable Aqua | Thermo Fisher | L34966 |
| KIR2DL1/S5 | KIR2DL1/S5 FITC | R&D Systems | FAB1844F |
| KIR3DL1 | KIR3DL1 PE | Biolegend | 312708 |
| CD45RA | CD45RA ECD | Beckman Coulter | B49193 |
| CD4 | CD4 PECy5.5 | Thermo Fisher | MHCD0418 |
| CD56 | CD56 PECy7 | Biolegend | 318318 |
| KIR2DL2/L3 | KIR2DL2/DL3 APC | Biolegend | 312612 |
| CD3 | CD3 APC Fire750 | Biolegend | 344840 |
| CD28 | CD28 BV421 | BD Horizon | 562613 |
| CD8 | CD8a BV711 | Biolegend | 301044 |
| CD16 | CD16 BV785 | Biolegend | 302046 |

**Table S14. Flow cytometry panel.**

### Supplementary text

#### Definition of Functional iKIR and inhibitory score.

An individual was defined to be positive for a functional KIR if they carried both the gene for the iKIR and the gene for its HLA ligand giving the following definitions:

| | |
|---|---|
| Functional *KIR2DL1* | *KIR2DL1* & *HLA-C2* |
| Strong Functional *KIR2DL2* | *KIR2DL2* & (*HLA-C1* or *B46* or *B73*) |
| Weak Functional *KIR2DL2* | *KIR2DL2* & *HLA-C2* |
| Functional *KIR2DL3* | *KIR2DL3* & (*HLA-C1* or *B46* or *B73*) |
| Functional *KIR3DL1* | *KIR3DL1* & *HLA-Bw4* |

Where *HLA-C1* denotes an HLA-C allele carrying a C1 motif (Asparagine at position 80); *HLA-C2* an HLA-C allele with a C2 motif (Lysine at position 80) and *HLA-Bw4* an HLA-B or HLA-A allele with a Bw4 motif (Asparagine at position 77). Despite carrying the Bw4 motif, HLA-A*25 does not bind to KIR3DL1 and so we do not consider *HLA-A*25* alleles as KIR ligands[36-38]. There is also evidence that, despite carrying a Bw4 motif, HLA-B*13 is not a KIR3DL1 ligand[36,38]. Exclusion of *HLA-B*13* alleles from the KIR ligand definition had no qualitative impact on the results.

Two metrics were considered: count and score. The count is the number of functional iKIR that an individual possessed, and the inhibitory score is the count adjusted for the observation that KIR2DL2 binds C1 more strongly than it binds C2 and that KIR2DL2 binds C1 more strongly than KIR2DL3 binds C1. Functionally diverse alleles at the same locus (2DL2/L3 and 3DL1/S1) were scored differently to reflect different strengths of inhibitory signal.

Inhibitory score= (1 if Func 2DL1) + (1 if Strong Func 2DL2 or 0.5 if weak Func 2DL2) + (0.75 if Func 2DL3) + (1 if Func 3DL1).

When it was necessary to stratify the cohort into individuals with a high count and low count or into individuals with a high inhibitory score and a low inhibitory score we analysed all thresholds that gave a sufficiently balanced stratification (similar number of people in each stratum).
